# Reversible fronto-occipitotemporal signaling complements task encoding and switching under ambiguous cues

**DOI:** 10.1101/2020.07.29.227736

**Authors:** Kaho Tsumura, Keita Kosugi, Yoshiki Hattori, Ryuta Aoki, Masaki Takeda, Junichi Chikazoe, Kiyoshi Nakahara, Koji Jimura

## Abstract

Adaptation to changing environments involves the appropriate extraction of environmental information to achieve a behavioral goal. It remains unclear how behavioral flexibility is guided under situations where the relevant behavior is ambiguous. Using functional brain mapping of machine-learning decoders and directional functional connectivity, we show that brain-wide reversible neural signaling underpins task encoding and behavioral flexibility in ambiguously changing environments. When relevant behavior is cued ambiguously during behavioral shifting, neural coding is attenuated in distributed cortical regions, but top-down signals from the prefrontal cortex complement the coding. When behavioral shifting is cued more explicitly, modality-specialized occipitotemporal regions implement distinct neural coding about relevant behavior, and bottom-up signals from the occipitotemporal region to the prefrontal cortex supplement the behavioral shift. These results suggest that our adaptation to an ever-changing world is orchestrated by the alternation of top-down and bottom-up signaling in the fronto-occipitotemporal circuit depending on the availability of environmental information.

## Introduction

Executive control guides flexible adaptation to changing environments, and is most developed in humans throughout evolution (Miller and Cohen 2001; Stoet and Snyder, 2009). Shifting between different types of behavior is one of the core executive control functions (Allport et al. 1994; Rogers and Monsell 1995), and task switching paradigms have been often used to investigate behavioral flexibility and its underlying neural mechanisms. Previous neuropsychological and neuroimaging studies of human and non-human animals suggest a critical role of the prefrontal cortex in switching tasks and rules (Dove et al. 2000; Rushworth et al. 2002; Bunge et al. 2005; Crone et al. 2006; Derrfuss et al. 2005; Kim et al. 2012; Yeung et al. 2006; Malagon-Vina et al. 2018; Bissonette et al. 2017; Fouragnan et al. 2019; Nee et al. 2016; Fig. 1A).

**Figure 1.**
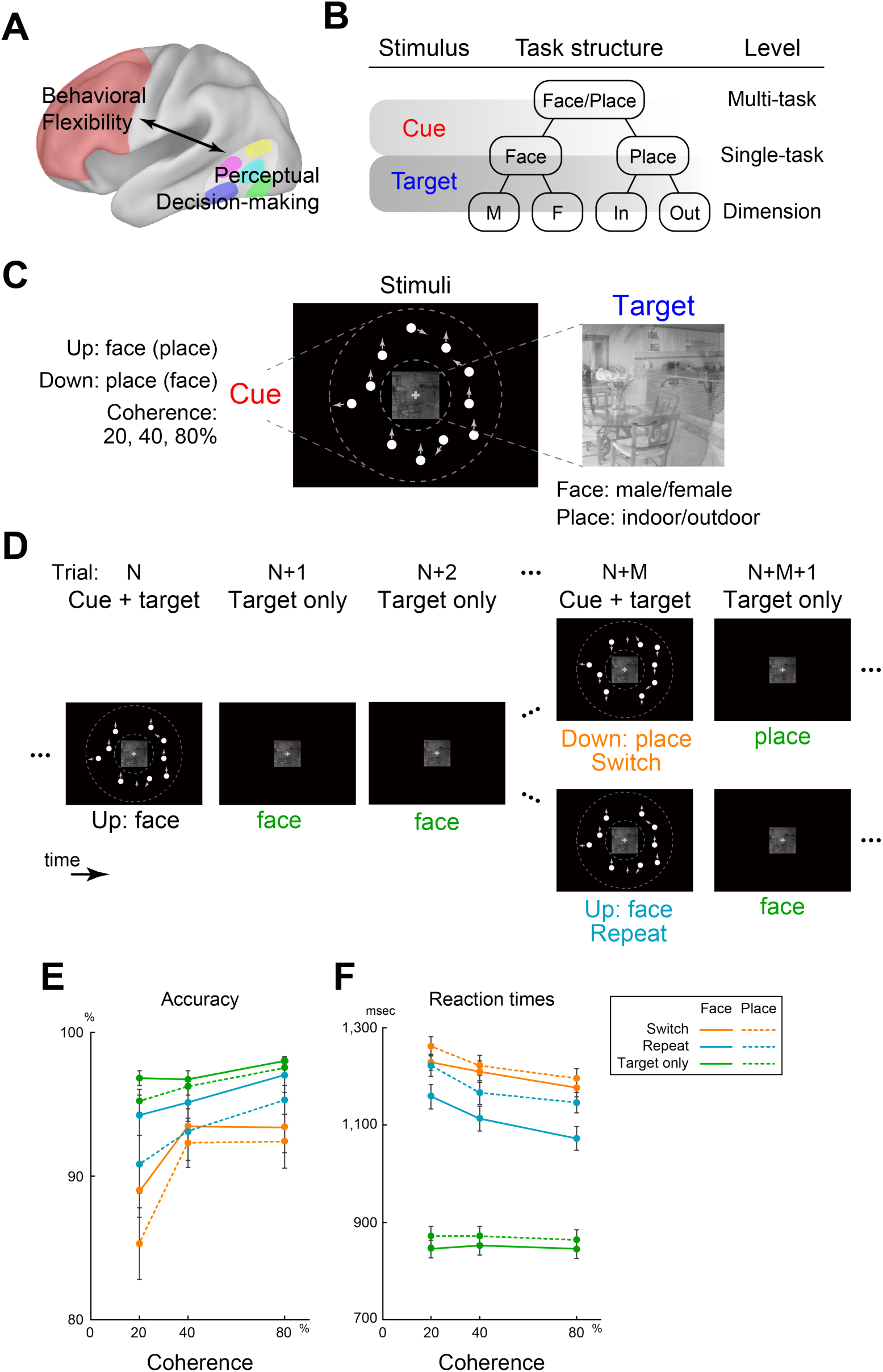
Experimental design and behavioral results. (A) Schematic illustration of a brain-wide model of interaction of behavioral flexibility and perceptual decision-making. Behavioral flexibility associated with prefrontal areas is subject to appropriate perception of the external world implemented in stimulus-modality-dependent occipitotemporal areas. (B) Hierarchical structure of task sets. Task sets are presented in a three-tiered decision tree. Behavioral flexibility is guided by the cue stimulus, which indicates relevant behavior (judging faces or places) and bridges the upper layers in the hierarchical structure. Base on the task to be performed, target stimulus is judged [male (M) or female (F) in the face task; indoor (In) or outdoor (Out) in the place task]. (C/D) Behavioral task. (C) Stimuli. Cue stimulus indicates the task to be performed (face or place task) and is composed of a set of white moving dots presented within a donut-shaped display circle (indicated by dotted lines). The arrow indicates the motion direction of each dot, and overall motion was either upward or downward. Upward and downward motion indicate face and place task, respectively. Motion strength of the cue stimulus was manipulated by coherence of dot motion. The target image was superimposed picture of face and place, which was presented at the center of screen. Participants judged whether the face picture was male or female, or the place is indoor or outdoor, depending on the task to be performed. (D) Task procedure. Trials with simultaneous presentation of a dot cue and target (switch and repeat; N and N+M th trials) were followed by target-only trials (N+1, N+2 and N+M+1 th trials). Switch and repeat trials were dependent on the performed task prior to the cue presentation. (E/F) Behavioral results. Accuracy (E) and reaction times (F) as a function of motion coherence. Solid and dotted lines indicate face and place task, respectively, and orange, blue, and green lines indicate switch, repeat, and target-only trials, respectively. Error bars indicate standard error of the mean across participants.

Importantly, executive control depends on perceived information of external environments, and relevant information is appropriately extracted from the external environment to achieve a behavioral goal. Perception of sensory information from the external environment guides the course of action, which is referred to as perceptual decision-making (Gold and Shadlen 2007; Hanks and Summerfield 2017). It involves the extraction of goal-relevant information, which is integrated to form a relevant decision. Studies of perceptual decision-making have used behavioral tasks that demand discrimination of sensory stimuli involving perceptual uncertainty (Newsome and Pare 1988; Corbetta et al. 1991; Kurikawa et al. 2018).

By manipulating perceptual uncertainty, neurophysiological and neuroimaging studies have examined cortical mechanisms of the perceptual decision-making. For example, the middle temporal (MT) region is known to play an important role in the perception of moving stimuli (Newsome and Pare 1988; Shadlen et al. 1996; Beauchamps et al. 1997, Huk et al. 2002; Kayser et al. 2010). It has been also suggested that the fusiform face area (FFA) and parahippocampal place area (PPA) are associated with the perception of face (Kanwisher et al. 1997; McCarthy et al. 1997; Ishai et al. 1999; Gazzaley et al. 2005; Freiwald and Tsao 2010) and place (Ishai et al. 1999; Gazzaley et al. 2005; Epistein and Kanwisher 1998) stimuli, respectively. These collective results suggest that temporal and occipital regions play important roles in perceptual decision-making, and are functionally segmented depending on the modality of the stimulus (Fig. 1A).

In our daily life, goal-relevant information in external environments is not always evident. Such situation can be well illustrated by an incorporation of behavioral shifting and perceptual decision-making. Prior studies have explored neural mechanisms during task switching under situations where target stimuli involved perceptual uncertainty (Kayser et al. 2010; Mante et al. 2013; Zhang et al. 2013; Kumano et al. 2016). More recently, we showed that increased uncertainty of target stimulus engaged top-down signals from prefrontal to occipitotemporal cortices, which complemented task switching in such situations (Tsumura et al. 2021).

Notably, the task switching paradigm is composed of hierarchically structured configuration of a set of task rules, called task sets (Bunge et al. 2005; Koechlin et al. 2003; Badre 2008; Jimura and Braver 2010; Fig. 1B). Those prior studies above manipulated perceptual uncertainty of the target items within the lower layers of the task set hierarchy (Kayser et al. 2010; Mante et al. 2013; Zhang et al. 2013; Kumano et al. 2016; Tsumura et al. 2021). In other words, the target stimulus to be discriminated was presented ambiguously, but the task *per se* was cued without ambiguity. To our knowledge, it remains unclear how task switch is achieved when the relevant task is indicated by a cue involving perceptual uncertainty. As such, the uncertainty of relevant task information in the upper layers of the task-set hierarchy may provide a novel and important opportunity to examine the relationships between executive control and perceptual decision-making (Figs. 1A/B).

One potential approach to elucidate underlying neural mechanisms is to identify the signal contents of responsible brain regions; this has recently been demonstrated for perception by neural decoding techniques (e.g. Kamitani and Tong 2005; Haynes and Rees 2006; Norman et al. 2006). In particular, prior neuroimaging studies have shown that mental and behavioral states can be decoded from neural coding by machine learning techniques that classify distributed patterns of brain activity (Kamitani and Tong 2005; Haynes and Rees 2006; Norman et al. 2006; Nishimoto et al. 2011; Loose et al. 2017; Qiao et al. 2017; Chikazoe et al. 2019; Wang et al. 2020). One commonly used technique is the support vector machine (SVM) (Vapnik 1998; Kamitani and Tong 2005; Misaki et al. 2010; Jimura and Poldrack 2012; Nakahara et al. 2016), which enables categorical discrimination by dividing multidimensional space composed of brain activation pattern using a linear hyperplane. Convolutional neural network (CNN) classifier (Krizhevsky et al. 2012; LeCun et al. 2015) is one of the deep neural network (DNN) classifiers consisting of multiple feature-aggregating layers, enabling more robust classification. Importantly, recent technical advancements of CNN allow mapping that highlights image locations characterizing a classified image (Selvaraju et al. 2017). The mapping technique may provide novel information about neural coding and functional localization of the brain.

The current study aimed to elucidate relationships between behavioral flexibility and perceptual decision-making under cue uncertainty, and to explore the underlying neural mechanisms (Figs. 1A/B). Functional MRI was administered while human participants performed a task-switching paradigm with a cue involving perceptual uncertainty. Standard univariate analysis identified brain regions associated with task switching, motion strength, and task modality. In order to elucidate causal network dynamics during task switching under cue ambiguity, we examined effective connectivity among the task-related brain regions. Finally, whole brain exploratory analyses based on machine learning techniques, CNN and SVM, were performed to identify brain regions that coded relevant task information.

## Materials and Methods

### Participants

Written informed consent was obtained from 30 healthy right-handed subjects (age range: 18-22; 11 females). Experimental procedures were approved by the institutional review board of Keio University and Kochi University of Technology. Participants received 2000 yen for each of the training and scanning sessions. One participant was excluded from analyses due to low behavioral performance; accuracy was lower than 30% in one of the experimental conditions. The number of participants was determined prior to the collection of the current data based on the effect sizes in pilot behavioral experiments and our previous relevant study (Tsumura et al. 2021).

### Behavioral procedures

The experiment consisted of two sessions administered on separate days. The first day was a training session, in which participants practiced discrimination tasks (random dot motion; Tsumura et al. 2021) and switching between two tasks (face and place tasks; see below for more details). On the second day, while fMRI scanning was administered, the participants performed the switching paradigm identical to those practiced in the training sessions.

### Stimuli

All stimuli were generated in Matlab version 2012a, using the Psychophysics Toolbox (Brainard 1997) extension version 3.0.10, and were visually presented on a computer screen. The current task cue stimuli were randomly moving dot stimuli similar to those used in previous studies of perceptual decision-making (Chen et al. 2015; Tsumura et al. 2021). Each motion stimulus involved 60 dots moving inside a donut-shaped display patch with a white cross in the center of the patch on a black background (Fig. 1C). The display patch and cross were centered on the screen and extended from 6 to 12^◦^ degrees of visual angle (dva). Within the display patch, every dot moved at the speed of 10 dva per second. Some dots moved coherently toward one direction (upward or downward) while the others moved randomly. The percentage of coherently moving dots determined the “motion coherence”, which was set to three levels (20, 40, and 80%).

Dot presentation was controlled to remove local motion signals following a standard method for generating motion stimuli (Newsome and Pare 1988; Britten et al. 1993; Palmer et al. 2005; Chen et al. 2015; Tsumura et al. 2021). Namely, upon stimulus onset, the dots were presented at new random locations on each of the first three frames. They were relocated after two subsequent frames, such that the dots in frame 1 were repositioned in frame 4, and the dots in frame 2 were repositioned in frame 5, and so on. When repositioned, each dot was either randomly presented at the new location or aligned with the pre-determined motion direction, depending on the pre-determined motion strength on that trial. Each stimulus was composed of 18 video frames with a 60 Hz refresh rates (i.e. 300-msec presentation).

Within the center circle mask of the donut-shape motion stimulus, a face/place superimposed stimulus was presented simultaneously (Fig. 1C). The face image set consisted of an image of picture of 4 male and 4 female unfamiliar Japanese faces, and the place image set consisted of 4 indoor and 4 outdoor unfamiliar places; this resulted in 64 overlaid images.

### Task procedure

At the beginning of the task, a dot patch and face/place stimulus were simultaneously presented. The direction of the dot patch (up or down) indicated the task to be performed (discrimination of face or place). Depending on the motion direction, participants were required to judge whether the face was male or female, or whether the place was indoor or outdoor (Fig. 1C), and pressed the corresponding button with their right thumb. Both of accuracy and speed were stressed. Stimulus was presented for 1.8 sec, followed by a 0.7 sec intertrial interval. If participants made an incorrect response, or did not respond within 1.8 sec from the stimulus onset, feedback stimulus indicating an error (X) was presented for 1.0 sec, followed by high-coherence (80%) cue trials for the same task dimension. The cue trials immediately after the error were discarded from analyses. Stimulus-response and cue-task associations for the two tasks were identical on days 1 and 2, and counterbalanced across participants.

The trial with simultaneous cue/target presentation was followed by trials with presentation of the face/place target stimulus without the dots cue stimulus (target only trials). The target-only trials were repeated for 3-5 times (Fig. 1D). In the target-only trials, participants were required to discriminate the center image stimulus along the same dimension until the next task cue (moving dots) was presented. One task block lasted for approximately for 90 secs, and 20-sec fixation blocks were inserted between task blocks. Each functional run involved 3 task blocks and lasted for 305 secs. The first trial at the beginning of each task block presented the dot cue with highest coherence (80%), and was discarded from analysis.

### Practice procedure

On the first day, participants practiced the tasks outside of the scanner. They first practiced a discrimination task for the moving dot stimulus, and were required to judge the direction of overall motion (up or down) and to press the correct corresponding button as quickly as possible (Tsumura et al. 2021). The response window was 1050 msec. Each practice run involved 70 trials, and 5 runs were administered for each participant. The first 5 trials and last 5 trials in each run used the highest coherence level (80%). Thus the middle 60 trials were composed of 20 trials for each of coherence levels (20, 40, or 80%).

The participants then practiced discrimination tasks for the face and place stimulus (see above). Across task switching practice runs, the switching frequency and coherence levels of the moving dot cue were manipulated such that the cue trials gradually became more difficult (i.e. more switch trials with low-coherence cue). Participants performed 8 practice runs for task switching.

### Behavioral procedure in scanning session

On the second day, after practicing task switching for one run, the participants performed 9 runs of task switching with identical procedure as for day 1 (see above) while functional MRI was administered. The frequency of switch and repeat trials and coherence level were approximately equivalent across runs.

### Imaging procedure

MRI scanning was administered by a 3T MRI scanner (Siemens Verio, Germany) with a 32ch head coil. Functional images were acquired using a multi-band acceleration echo-planar imaging sequence (Moeller et al. 2010) [repetition time (TR): 0.8 sec; echo time (TE): 30 msec; Flip angle (FA): 45 deg; 80 slices; slice thickness: 2 mm; in-plane resolution: 3 x 3 mm; multiband factor: 8]. Each functional run involved 385 volume acquisitions. The first 10 volumes were discarded from analysis to take into account the equilibrium of the longitudinal magnetization. High-resolution anatomical images were acquired using an MP-RAGE T1-weighted sequence [TR: 2500 msec; TE = 4.32 msec; FA: 8 deg; 192 slices; slice thickness: 1 mm; in-plane resolution: 0.9 x 0.9 mm^2^].

### Behavioral analysis

Trials were classified into three types in terms of switching: 1) task switch trials presenting the random dot cue and target stimuli simultaneously, where the task to be performed alternated (i.e., face to place or place to face); 2) task repeat trials presenting the random dot cue and target stimuli simultaneously, where the same task was repeated; and 3) target-only trials presented after the switch and repeat trials, and their subsequent trials without random dot cue stimuli presentation (Fig. 1D). These trial types were analyzed separately. Trials were also classified by task dimension (face or place), and switch and repeat trials were examined at each coherence level (20, 40, or 80%). Accuracy and reaction times (RTs) were calculated for each trial condition, and then compared. Statistical testing was performed based on repeated measures ANOVAs implemented in SPSS Statistics 24 (IBM Corporation, NY USA).

### Image preprocessing

MRI data were analyzed using SPM12 software (http://fil.ion.ac.uk/spm/). All functional images were initially temporally realigned across volumes and runs, and the anatomical image was coregistered to a mean image of the functional images. The functional images were subsequently spatially normalized to a standard MNI template with normalization parameters estimated based on the anatomical scans. The images were resampled into 2-mm isotropic voxels, and spatially smoothed with a 6-mm full-width at half-maximum (FWHM) Gaussian kernel.

### Imaging analysis: general linear model (GLM)

#### Single level analysis

A GLM approach (Worsley and Friston 1995) was used to estimate parameter values for task events. The events of interest were correct switch, repeat, and target-only trials. For switch and repeat trials, the normalized (z-scored) coherence level of the dot stimuli was also added as a parametrical effect of interest (Tsumura et al. 2021). Error trials in all conditions were separately coded in GLM as nuisance effects. Those task events were time-locked to the onset of target images and then convolved with canonical hemodynamic response function (HRF) implemented in SPM. Additionally, six-axis head movement parameters, white-matter signals, lateral-ventricle signals, and parametrical effect of reaction times normalized across trials were also included in GLM as nuisance effects. The parameters were then estimated for each voxel across the whole brain.

#### Group-level analysis

Maps of parameter estimates were first contrasted within individual participants. Contrast maps were collected from all participants, and subjected to a group-level paired t-test. For the coherence effect, the contrast maps were subjected to a one-sample group-mean test, with maps weighted and summed based on normalized coherence levels. Voxel clusters were identified using an uncorrected threshold of P < .001 based on voxel-wise t-statistics. The voxel clusters were tested for a significance with a threshold of P < .05 corrected by family-wise error (FWE) rate based on permutation methods (Nichols and Holmes 2001) (5000 permutations) implemented in *randomise* in FSL suite (http://fmrib.ox.ac.uk/fsl/). This group analysis procedure was validated to appropriately control false positive rates in a prior study (Eklund et al. 2016). Peaks of significant clusters were then identified and listed on tables. If multiple peaks were identified within 12 mm in one cluster, the most significant peak was retained. When exploring brain regions associated with motion coherence, exploration was restricted within a mask obtained from Neurosynth (Yarkoni et al. 2011) (http://neurosynth.org/) for the search word ‘motion’ (z > 3.0, for uniformity test), in order to ensure the extraction of motion-related regions, because the current cue trials simultaneously presented face/place stimuli in addition to cues indicating task switching tasks between face and place discrimination.

#### Effective connectivity analysis

The current analysis was designed to test the hypothesis that functional connectivity among brain regions associated with task switching, motion perception, face perception, and place perception identified in univariate analysis (Figs. 2A-C) is modulated by task manipulations and brain signals. Dynamic causal modeling (DCM; Friston et al. 2003) analysis was performed in order to examine functional connectivity mechanisms associated with task switching under cue uncertainty (Figs. 2D-E). DCM allows us to explore the effective connectivity among brain regions under the premise that the brain is a deterministic dynamic system that is subject to environmental inputs and produces outputs based on the space-state model. The model constructs a nonlinear system involving intrinsic connectivity, task-induced connectivity, and extrinsic inputs. Parameters of the nonlinear system are estimated based on fMRI signals (system states) and task events. The use of a high temporal resolution sequence for functional imaging enables us to collect more scan frames to increase the signal-to-noise ration of the DCM analysis (Penny et al. 2004; Stephan et al. 2010; Tsumura et al. 2021).

Four regions of interest were first defined based on univariate analysis and prior studies of task switching and perceptual decision-making: 1) task switching [left lateral prefrontal cortex (lPFC) (Dove et al. 2000; Konishi et al. 2002; Rushworth et al. 2002; Bunge et al. 2005; Crone et al. 2006; Derrfuss et al. 2005; Yeung et al. 2006; Kim et al. 2012; Jimura and Braver 2010; Tsumura et al. 2021); Fig. 2A, see also Results]; 2) motion perception [middle temporal (MT) (Gold and Shadlen 2007; Hanks and Summerfield 2017; Newsome and pare 1988; Shadlen et al. 1996; Beauchamps et al. 1997, Huk et al. 2002; Kayser et al. 2010; Tsumura et al. 2021); Fig. 2B, see also Results]; 3) face perception [fusiform face area (FFA) (Kanwisher et al. 1997; McCarthy et al. 1997; Ishai et al. 1999; Gazzaley et al. 2005; Freiwald and Tsao 2010); Fig. 2C, see also Results]; 4) place perception [parahippocampal place area (PPA) (Ishai et al. 1999; Gazzaley et al. 2005; Epistein and Kanwisher 1998); Fig. 2C, see also Results]. More specifically, meta-analysis maps were obtained from Neurosynth (http://neurosynth.org/; Yarkoni et al. 2011) using a keyword search for “switching”, “mt”, “ffa”, “place” to obtain the meta-analysis maps for lPFC, MT, FFA, and PPA regions of interest (ROIs), respectively. ROI images were then created with 6-mm radius spheres centered in the peak coordinates in the meta-analysis activation maps thresholded above z > 10 (uniformity test).

**Figure 2.**
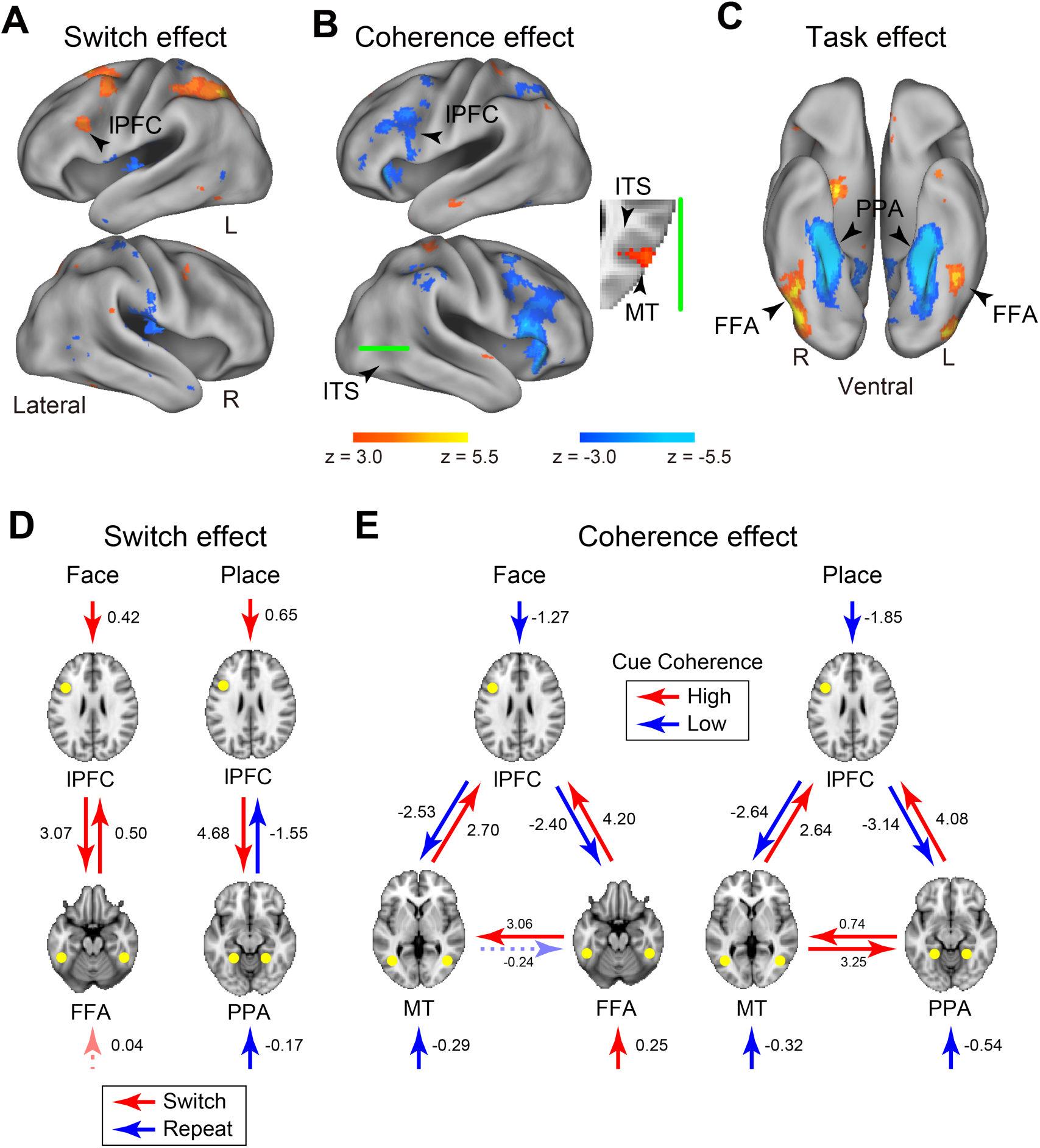
Whole-brain exploration activation and functional connectivity analyses (A-C). Statistical activation map of univariate analysis and functional connectivity analysis. Maps are overlaid onto the 3D surface of the brain. Hot and cool colors indicate positive and negative effects, respectively. (A) Switch effect (switch minus repeat trials) (B) Motion coherence effect (high- vs. low-coherence trials). Green solid line on the surface indicates the axial section on the right. (C) Task effect (face minus place tasks in target only trials). lPFC: lateral prefrontal cortex; ITS: inferior temporal sulcus; MT: middle temporal; FFA: fusiform face area; PPA: parahippocampal place area. (D/E) Effective connectivity analysis. (D) Effective connectivity and extrinsic inputs modulated by the contrast switch vs. repeat trials during face (left) and place (right) tasks. Red and blue arrows indicate connectivity enhancements in switch and repeat trials, respectively. The values next to the arrows indicate the magnitude of connectivity enhancements (positive: switch > repeat; negative: repeat > positive). (E) Effective connectivity and extrinsic inputs modulated by motion coherence during face (left) and place (right) tasks. Red and blue arrows indicate enhancements in high- and low-coherence trials, respectively. The values next to the arrows indicate the magnitude of connectivity (positive: high coherence > low coherence; negative: low coherence > high coherence). Arrows with solid line indicate statistically significant connectivity (P < .05, uncorrected).

Given these ROIs, we first tested whether the switch-related prefrontal region sends (or receives) a task-related signal toward (or from) the stimulus-modality dependent occipitotemporal region of the target (i.e. FFA/PPA) during task switching (Fig. 2D). If this were the case, then we tested whether signaling between prefrontal and occipitotemporal regions changed depending on the uncertainty of cue stimuli (Fig. 2E).

Signal time courses of four ROIs and regressors in events of interest were extracted from first-level GLMs. The events of interest were cue (switch and repeat) trials and target-only trials of each task. For switch and repeat trials, the contrast of the two trials and normalized coherence level of the dot stimuli were added as parametrical effects of interest. Nuisance effects of head-motion, white matter signal, ventricle signal, functional run, and contrast were subtracted out from the ROI timecourses. The input matrix was U mean-centered.

For each trial effect, causal models were defined as those that differed in external inputs and modulatory effects among ROIs. Because the current analysis involved 2 or 3 ROIs (Figs. 2D-E), the tested models included 16 or 512 types (i.e. 2^2^ inputs and 2^2^ connection effects or 2^3^ inputs and 2^6^ connection effects). Connectivity matrices reflecting 1) first-order connectivity, 2) effective change in coupling induced by the inputs, and 3) extrinsic of inputs on MRI signal in ROIs were estimated for each of 1the 6 or 512 models based on DCM analysis implemented in SPM12. Parametric regressor (switch vs repeat / coherence) was used as an extrinsic effect for effective connectivity between ROIs and ROIs inputs.

In order to estimate strength of effective connectivity, a Bayesian model reduction method (Friston et al. 2016) was used. The reduction method allows the calculation of posterior densities for all possible reduced models, which were then inverted to a fully connected model. The reduced models were then supplemented by second-level parametric empirical Bayes (Friston et al. 2016) (PEB) to apply empirical priors that remove subjects’ variability from each model.

Next, the parameters of these models were estimated based on Bayesian model averaging (Friston et al. 2003) (BMA) to estimate group-level statistics. Because the current analysis aimed to identify effective connectivity observed as an average across participants, we used a fixed effect (FFX) estimation assuming that every participant uses the same model. This is in contrast to using a random effect (RFX) estimation assuming different participants use different models, which is often used to test group differences in effective connectivity (Penny et al. 2010). The significance of connectivity was then tested by thresholding at a posterior probability at the 95% confidence interval. We used the uncorrected threshold, because the current analysis aimed to test if connectivity between two specific brain regions was enhanced depending on task manipulation and brain activity, not to explore one model involving connectivity among multiple brain regions that best fits to the imaging and behavioral data (Tsumura et al. 2021).

Additionally, in order to test the robustness of the functional connectivity, we performed supplemental analyses. We estimated model parameters 1) without an empirical prior (Figs. S1C-D); 2) changing the number of the regions of interest (ROIs) in the models (Figs. S1E-F), and 3) changing the definition of the ROIs (Figs. S1G-H). When changing the ROI definition, we used a leave-one-out procedure; the centers of ROIs of one participant were determined based on group-level univariate activation maps of corresponding contrasts (i.e. lPFC: switch vs. repeat trials; MT: high vs. low coherence trials; FFA/PPA: face vs. place of target-only trial) without the participant in order to circumvent circular analysis. The ROIs were created as spheres with 6-mm radius for individual participants. Complete results can be provided upon request.

### Convolutional neural network (CNN) classifier

In order to explore brain regions involving task-related neural representation, a convolutional neural network classifier (CNN) (Krizhevsky et al. 2012; LeCun et al. 2015) was used. The current CNN model was based on VGG16 (Simonyan et al. 2015), with five convolution layers for extracting image features and two fully connected layers for binary classification. Initial parameters of convolution layers were set to parameters pre-trained with concrete object images provided from ImageNet (http://www.image-net.org/; Deng et al. 2009; Fig. S2A).

The VGG16/ImageNet model is capable of classifying concrete object images into 1,000 item categories. Importantly, it has been demonstrated that the pre-trained model can learn novel image sets more efficiently than the non-trained model by tuning convolution and fully connected layers and fully connected layers only (Donahue et al. 2014; Pan et al. 2010; Fig. S2C). Thus, the current analysis retrained the pre-trained VGG16-ImageNet model to classify brain activation maps.

Training data were single-subject 2nd level z-maps during the N-back working memory task from the S1200 release of the Human Connectome Project (N = 992; HCP; http://www.humanconnectomeproject.org/; Barch et al. 2013; Glasser et al. 2016a). From each participant, statistical z-maps for activation contrasts for face vs. fixation and place vs. fixation (2-back and 0-back corrupted) were collected. We used gray-scaled flat 2D cortical maps (Glasser et al. 2016b) provided from HCP (992 images; face: 496, place: 496; Fig. S2B) for dimensional compatibility of images between VGG16-ImageNet and activation maps. The training data set was divided into 10 subsets, and 9 subsets were used for retraining and the remaining 1 set was used for validation, enabling a 10-fold cross-validation test. Then, the pre-trained VGG16 model was retrained by the activation maps such that the model classifies face and place trials. Model training and testing was implemented using Keras (https://keras.io/) under Tensorflow backend (https://www.tensorflow.org/) [input image size: 480 x 1280 pixels; batch size: 10; epoch: 50; learning rate: 0.0001; optimizer: Stochastic Gradient Descent (SGD); Fig. S2C *top*].

After retraining of the HCP working memory maps, the model with highest classification accuracy was further retrained to classify activity maps for face and place tasks during target-only trials of the current dataset (Fig. S2C *bottom*). For each functional run of each participant, a single-level GLM estimation was performed with regressors identical to those in the univariate analysis as described above. The GLM estimations were performed within standard MNI space. Activation z-maps for the contrasts for face vs. fixation and place vs. fixation during correct target-only trials were collected from each functional run. Activity maps for the contrast for face vs. fixation and place vs. fixation were then gray-scaled and flattened such that these maps were anatomically and geometrically identical to those from the HCP working memory task using Connectome Workbench (https://www.humanconnectome.org/software/connectome-workbench/). The training dataset consisting of 522 images (261 face and 261 place maps from each of 9 runs of 29 participants) was divided into 10 subsets, enabling a 10-fold cross-validation test.

Given the limited number of images available from the current experiment, this two-step retraining of the model was found to be effective when classifying current tasks, because 1) training randomly initialized models failed in classifying the current target-only trials (Fig. S3A) and in classifying the HCP working memory conditions (Fig. S3B); 2) retraining VGG16-ImageNet was successful in classifying the HCP working memory conditions (Fig. S3B); 3) retraining VGG16-ImageNet model failed in classifying the current target-only trials (Fig. S3A); and 4) retraining VGG16-ImageNet/HCP was successful when classifying the current face/place tasks (Fig. S3A).

After learning of the target-only trials from the current dataset, the retrained 10 models were tested to classify activation maps during task switching and repeat trials where the dot cue stimulus was presented to indicate the task to be performed. Testing data was created based on a GLM analysis where switch and repeat trials at differential coherence levels were coded separately. For each functional run of each participant, a single-level GLM estimation was performed, and activation contrast z-maps for face vs. fixation and place vs. fixation were collected during those correct switch and repeat trials. Gray-scale 2D activation maps were created for the contrasts for face vs. fixation and place vs. fixation during 6 types of cue presentation trials (switch/repeat x high/middle/low coherence). Importantly, the testing data were independent of the 2 sets of retraining data (HCP working memory and current target-only trials). The maps were tested, and accuracy was averaged across cross-validation models within each participant. A statistical test of classification accuracy was performed based on repeated measures ANOVAs implemented in SPSS Statistics 24 (IBM Corporation, NY USA).

In a separate supplemental analysis, in order to examine whether the current results were biased by subject-specific characteristics of image data, we used a leave-one-subject-out procedure to retrain the CNN classifier to classify activation maps of the target only trials, and then tested the remaining subject (Fig. S5, see also Results).

A recent study demonstrated that a CNN model successfully learned and classified task-related fMRI images without flattening the images (Wang et al. 2020). Although retraining of the classifier was also effective for small data sets, this technique is available only for blocked-design experiments as the model was trained and tested based on fMRI timeseries, which increased the number of the training data. The model also normalizes and convolves the timeseries along the temporal axis. Because the current study used event-related design, only activation maps estimated by single-level GLM analyses were available for training and testing. Because of the nature of GLM analysis, the number of available images were limited in comparison with fMRI timeseries. Thus, retraining of pre-trained model based on flattened 2D activation maps is effective for small size dataset of event-related fMRI.

### Visualization of convolution layer weights

In order to identify brain regions involving discriminative information to classify performed tasks, Grad-CAM (Selvaraju et al. 2017) was used. Grad-CAM visualizes the aggregation of weight gradients between convolution layers, and highlights image locations critical for classification on a pixel-by-pixel basis (Fig. 3C).

**Figure 3.**
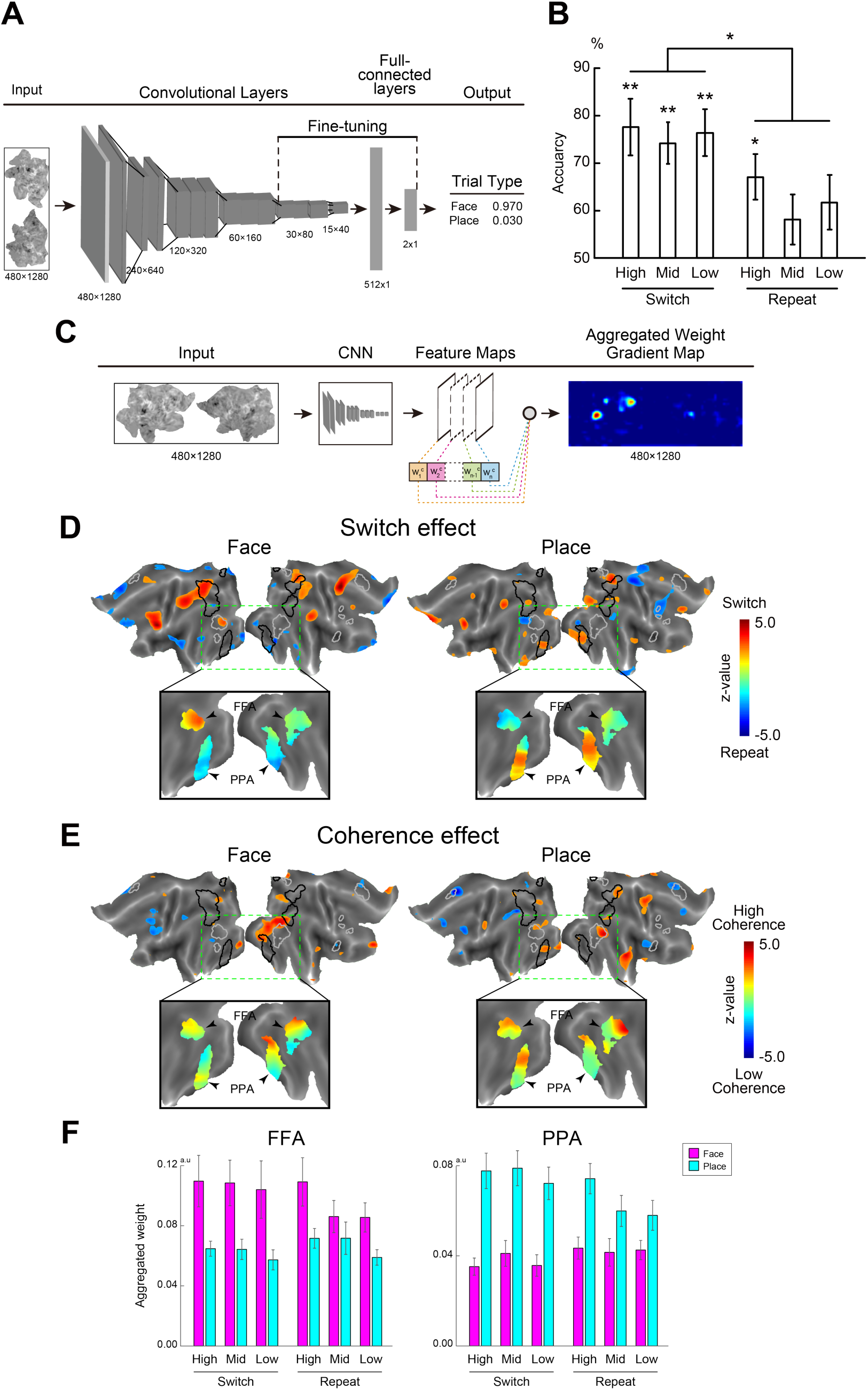
Convolutional neural network (CNN) classifier mapping. (A) CNN model was based on VGG16 pre-trained by ImageNet data. The model was retrained to classify performed task (face or place). Retraining of brain maps was based on fine tuning from the 5th convolution layer to the full-connected layers. (B) Classification accuracy for each task condition. *: P < .005; **: P < .001. High: high-coherence (80%) trials; Mid: middle-coherence (40%) trials; Low: low-coherence (20%) trials. (C) Weight gradients between convolution layers was aggregated and then visualized to identify cortical regions involving more information to classify tasks. (D) Visualization of aggregated weight contrasts for switch vs. repeat trials in face task (left) and place task (right). Statistical maps are overlaid onto flat cortical anatomical images with a statistical threshold of |z| > 2.0 (top). Hot and cool colors indicate statistical z-values of positive (switch > repeat) and negative (repeat > switch) effect, respectively. Gray and black closed lines overlaid on flat map indicate brain regions significantly activated during face and place tasks in univariate analysis, respectively (Fig 2C and S6 Fig). Occipitotemporal regions in rectangular boxes with green broken lines were expanded below. Maps were overlaid onto flat maps masked by the univariate activation contrast face vs. place tasks for target only trials (gray and black closed lines in the top panels). The fusiform face area (FFA) and parahippocampal place area (PPA) are indicated by arrow heads. (E) Visualization of weight contrast for coherence effect in face task (left) and place task (right). Hot and cool colors indicate greater statistical values of weights in high- and low-coherence trials, respectively. The formats are similar to those in panel (D). (F) Regions of interests (ROIs) analysis. PPA and FFA ROIs were defined based on target-only trials, and weight magnitudes were collected for each cue trial conditions and ROIs. All error bars in the figure indicate standard errors of the means across participants.

Aggregated weights were visualized onto 2D brain surface maps for each tested map. For each participant, the weight maps were averaged across models and maps within each of the 6 cue trial conditions (switch/repeat x high/middle/low coherence cue). The averaged maps were contrasted between switch and repeat trials to explore brain regions showing differential weights gradients between switch and repeat trials within participants. For coherence the effect, three maps for coherence level trials were weighted and summed based on behavioral accuracy estimated by sigmoid fitting within participants. For each contrast, maps were collected from all participants, and pixel-wise z-values were calculated treating participants as a random effect, and the z-values were mapped on the 2D surfaces of the cortical areas.

In order to statistically test dissociable weight patterns during cue trials in the FFA and PPA, ROI analyses were performed (Fig. 3F). ROIs were defined as occipitotemporal regions showing greater activation in the contrast of face minus place tasks or place minus face tasks (Figs. 2C and S4; Table S3), independently of the cue trials. From each ROI, weight magnitudes were collected for each of the trial conditions: task (face/place), switching (switch/repeat), and cue coherence [high/middle/low: 80/40/20%], and then averaged within ROIs for each participant. Statistical tests were then performed based on repeated measures ANOVA.

### Support vector machine (SVM) analysis for whole brain cortical regions

In order to supplement the CNN classifier analysis, multi-variate pattern analyses (MVPA) based on SVM were performed. The SVM classifier was trained to perform bivariate classification for face and place tasks. Training and testing were implemented using scikit-learn package (https://scikit-learn.org/stable/) with a Tensorflow backend (https://www.tensorflow.org/). We used a linear kernel, and adjusted C parameters (C = 0.1, 1.0, 10.0). As the overall results were maintained with the C adjustment, then we reported the results with the default parameter (C = 1.0).

For classifier training, we used the image set of single-subject 2nd level z-maps during the N-back working memory task (face vs. fixation and place vs. fixation) obtained from the S1200 HCP (N = 992), which was identical to those used in the CNN classifier training. The classifier was trained by the activation images, such that it classified face and place tasks during the HCP N-back working memory task. Weights of the trained classifier were mapped on the 2D cortical surface.

The testing dataset was also identical to those used in the CNN classifier analysis: 2-D z-maps for activation contrast of switch and repeat trials (switch vs. fixation and repeat vs. fixation) at each coherence level (20, 40, 80%) of the current experiment (Fig. S2B). Testing activation images were subject to the trained classifier for each cue trial condition of each participant, and classification accuracy was averaged for each trial condition across participants (Fig. 4A). In a separate analysis, the SVM classifier was trained based on single level z-maps during target only trials from the current dataset (N = 29), and the identical image set was tested (Fig. S6).

**Figure 4.**
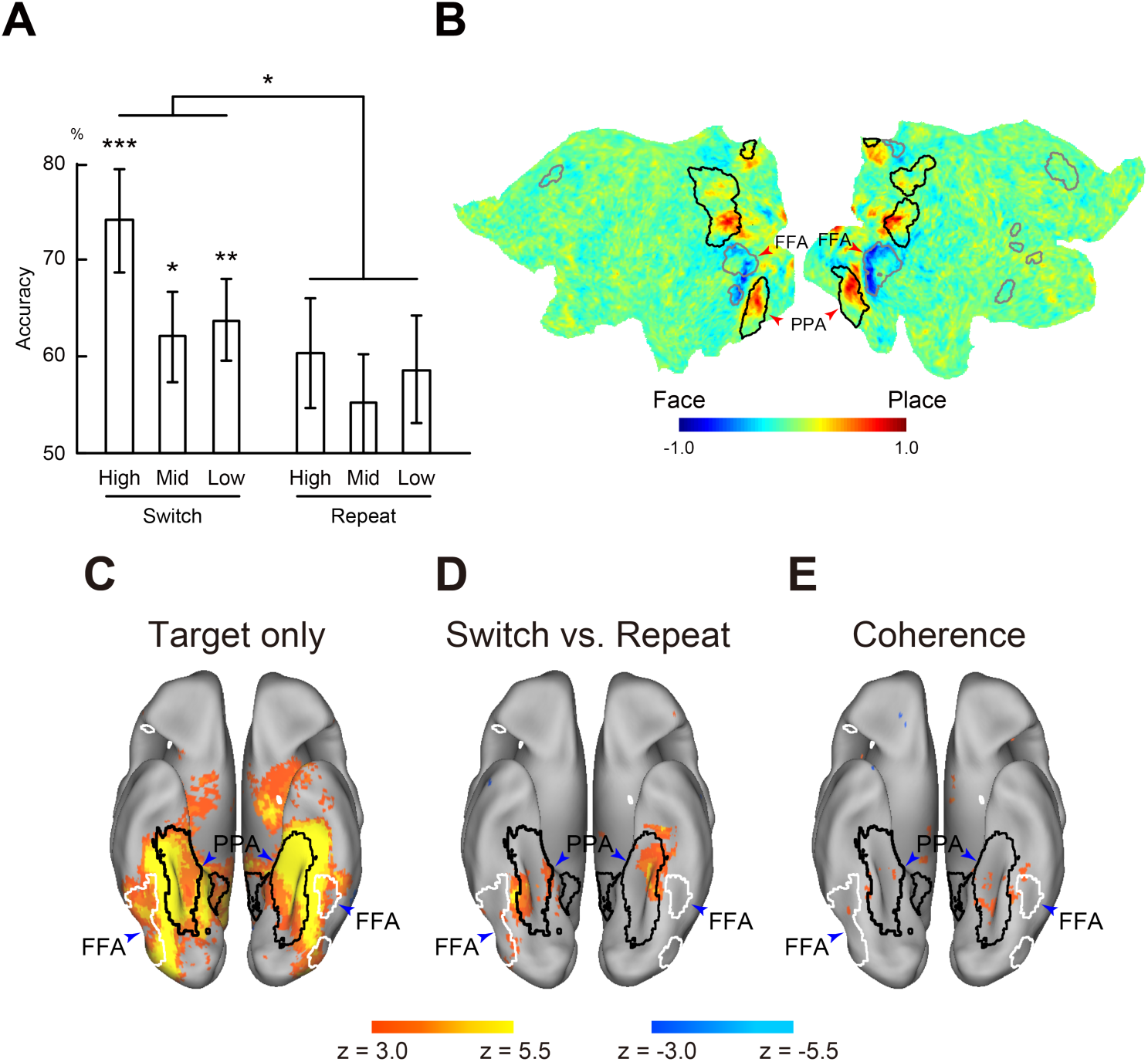
Multi-variate pattern analysis (MVPA) with support vector machine (SVM). (A) Classification accuracy for each task condition with SVM classification in the whole brain. The model was trained for classification of face and place tasks with the working memory task in the Human Connectome Project (HCP) and tested with each condition of cue trials in the current study. Error bars indicate standard error of the mean across participants. *: P < .05; **: P < .005; ***: P < .001. (B) Visualization of weight assigned to the pixels for classification of face and place tasks in the working memory task in the HCP. Formats are similar to those in Figs. 3D/E. (C-E) Statistical significance maps for searchlight MVPA. Classifiers were trained to classify the performed task. Maps are overlaid onto a 3D surface of the brain and displayed from a ventral view. White and black closed lines overlaid onto the 3D surface of the brain indicate significant clusters for contrast face vs. place tasks in univariate analysis, respectively (Fig. 2c). The fusiform face area (FFA) and parahippocampal place area (PPA) are indicated by blue arrow heads. (C) Target-only trial. Hot and cool colors indicate statistical level for classification accuracy relative to chance level. (D) Accuracy difference between switch and repeat trials. Hot and cool colors indicate higher accuracy in switch and repeat trials, respectively. (E) Differential classification accuracy depending on the coherence effect. Hot and cool colors indicate higher accuracy in high- and low-coherence trials, respectively.

### Searchlight SVM

In order to explore brain regions in which local activity patterns involves information about a performed task (face/place), searchlight MVPA (Kriegeskorte et al. 2006) was conducted. Bivariate classification based on SVM was used to decode the performed task (face or place). A searchlight procedure with a 5-voxel radius was used to provide a measure of decoding accuracy in the neighborhood of each voxel. Training and testing were performed based on the Decoding Toolbox (TDT; version 3.95; https://sites.google.com/site/tdtdecodingtoolbox/). Again, training and testing data were independent, based on different behavioral tasks and data sets (training: HCP working memory; testing: current task switching with male/female or indoor/outdoor judgements).

Training data were 3D single subject 2nd level z-maps during N-back working memory task (N = 1,000) from HCP S1200 release (Barch et al. 2013; Glasser etl al. 2016a). Similar to the CNN analysis, we used activation contrasts for face task vs. fixation and for place task vs. fixation (2-back and 0-back corrupted). These activation contrasts were collected from each participant, and the whole dataset was divided into 10 subsets (N = 100 each). For each subset of the training data, a classifier in each searchlight was trained based on these z-maps such that it classified face and place conditions during the HCP N-back working memory task.

Test data were single-subject z-maps during switch, repeat and target only trials of the current experiment. For each functional run of each participant, single-level GLM estimation was performed with regressors identical to those in the univariate analysis as described above. Activation maps for face vs. fixation and place vs. fixation during switch and repeat correct trials were collected from each functional run. These GLM analyses were performed within standard MNI space.

Another set of testing data was created based on a separate GLM analysis with correct cue (switch and repeat) trials separately coded at each coherence level (20/40/80%). For each functional run of each participant, a single-level GLM estimation was performed, and activation maps for face vs. fixation and place vs. fixation were collected for each coherence level.

For each training subset, the classifier was tested on whether it correctly classified the performed task (face or place task) for each trial condition (i.e. switch/repeat/target only and coherence level). Classification performance was then collected from all functional runs and averaged within participants for each searchlight. The performance of classification was calculated as the accuracy minus chance level for bivariate classification. Accuracy maps were then averaged across testing data sets within participants.

Accuracy maps for switch, repeat and target only trials were first averaged across training subset models within participants, and averaged accuracy maps were collected from all participants. Voxel-wise one-sample group-mean test was performed for each trial condition, with a procedure similar to that in the univariate analysis as stated above. In order to explore brain regions showing differential classification accuracy between switch and repeat trials, voxel-wise group-level paired test was performed, and significance was tested similarly.

Accuracy maps for each coherence level trials were also averaged across training subset models within participants, and the averaged accuracy maps were collected from all participants. Voxel-wise one-sample group-mean test was performed for each coherence level, and significance was tested similarly. In order to examine coherence effect, the maps for three coherence levels were weighted and summed based on behavioral accuracy estimated by sigmoid fitting. Voxel-wise one-sample group-mean test was performed, and significance was tested similarly.

In separate analyses, all group-level tests above were also performed for each of the 10 training subset models, and we confirmed that overall results were consistent.

## Results

### Behavioral results

Human participants performed a task-switching paradigm (Bunge et al. 2005; Koechlin et al. 2003; Badre 2008; Jimura and Braver 2010; Tsumura et al. 2021), in which they alternated discrimination tasks for face and place stimuli (Figs. 1C/D). The relevant task was indicated by a cue stimulus involving perceptual uncertainty, which was manipulated by the motion strength of randomly moving dots.

Accuracy was lower in low-coherence (i.e., more uncertain) trials compared to high-coherence trials [F(1, 28) = 12.7; P < .005; Fig. 1E], and became lower in switch trials than in repeat trials [F(1, 28) = 11.8; P < .005]. Likewise, reaction times (RTs) were longer in low-coherence trials compared to high-coherence trials [F(1, 28) = 69.9; P < .001; Fig. 1F], and were longer in switch trials than in repeat trials [F(1, 28) = 43.2; P < .001]. These behavioral results suggest that the current behavioral task successfully manipulated task switching (Allport et al. 1994; Rogers and Monsell 1995; Dove et al. 2000; Rushworth et al. 2002; Crone et al. 2006; Yeung et al. 2006; Jimura and Braver 2010; Tsumura et al. 2021) and perceptual decision-making (Newsome and Pare 1988; Shadlen et al. 1996; Palmer et al. 2005; Kayser et al. 2010; Hanks and Summerfield 2017; Tsumura et al. 2021). The interaction effect of trial type (switch/repeat) and coherence levels did not reach statistical significance [F(1,28) = 2.1; P = 0.155].

In trials without cue presentation, occurring after the switch and repeat trials until the next cue trials (target only trial; Fig. 1D), accuracy was lower after lower coherence cue trials [F(1, 28) = 16.3; P <. 001; Fig 1E], but RTs remained unchanged after lower coherence cue trials [F(1, 28) = 1.0; P = 0.317], suggesting that switching might not complete immediately after low coherence cue trials. RTs were longer in place than face tasks [F(1, 28) = 5.0; P <. 05; Fig. 1F].

### Exploration of switch-related and stimulus-modality-dependent brain regions

We first explored brain regions associated with task switching, motion coherence, and the perception of face and place based on univariate general linear model (GLM) analysis. Figure 2A shows brain regions showing significant increases and decreases in univariate brain activity during switch relative to repeat trials (P < .05 corrected with cluster-wise family-wise error rate based on non-parametric permutation tests; see Materials and Methods). Robust activation increases were observed in the left frontal regions including the inferior frontal cortex (IFC), dorsolateral prefrontal cortex (DLPFC), inferior frontal junction (IFJ), and pre-supplementary motor area (pre-SMA), and in left parietal regions including posterior parietal cortex (PPC), consistent with prior studies (Dove et al. 2000; Konishi et al. 2002; Rushworth et al. 2002; Bunge et al. 2005; Crone et al. 2006; Derrfuss et al. 2005; Yeung et al. 2006; Kim et al. 2012; Jimura and Braver 2010; Tsumura et al. 2021). A full list of brain regions is shown in Table S1.

We then explored brain regions associated with motion coherence. Figure 2B shows brain regions showing significant modulation of brain activity in relation to motion coherence during the cue (i.e. switch and repeat) trials. In low-coherence trials, activation was increased in multiple fronto-parietal regions including IFC, DLPFC, IFJ, pre-SMA, and PPC (Fig. 2B and Table S2), consistent with prior studies (Kayser et al. 2010; Tsumura et al. 2021). In contrast, activation was greater in high-coherence trials in the MT region (Fig 2B *right*; Table S2), which is also consistent with prior studies of perceptual decision-making for motion (Newsome and Pare 1988; Shadlen et al. 1996; Beauchamp et al. 1997; Huk et al. 2002; Kayser et al. 2010; Tsumura et al. 2021).

We next explored brain regions associated with face and place tasks (Fig. 2C and Table S3). Consistent with prior studies of perception of face (Kanwisher et al. 1997; McCarthy et al. 1997; Ishai et al. 1999; Gazzaley et al. 2005; Freiwald and Tsao 2010) and place (Ishai et al. 1999; Gazzaley et al. 2005; Epistein and Kanwisher 1998), in the face task, activity was greater in the FFA, whereas in the place task, increased activity was observed in the PPA.

These collective univariate activation results suggest that the current univariate activation analysis successfully identified brain regions associated with task switching in addition to the perception of face, place and motion stimulus, and that those regions were cooperatively engaged in task switching when the cue stimulus involved perceptual uncertainty.

### Reversal of functional connectivity depending on cue uncertainty

The whole brain exploratory analyses of univariate activation identified three types of brain regions: 1) left PFC associated with task switching and perceptual uncertainty (Figs. 2A-B); 2) MT region associated with motion coherence of task cue (Fig. 2B); and 3) FFA and PPA associated with discrimination of face and place stimuli, respectively (Fig. 2C). One possible mechanism to explain these regions playing differential roles in task switching with cue uncertainty is that lPFC, MT, and FFA/PPA mutually received or sent task-related signals during switching, which was modulated by cue coherence, such that the task-related signal complemented the engagement of these regions depending on cue uncertainty and the task to be performed.

In order to test this hypothesis, we performed an interregional effective connectivity analysis based on dynamic causal modeling (DCM) that allows the examination of directionality of task-related functional connectivity based on the state-space model (see Materials and Methods).

We first examined the task-related effective connectivity during task switching, and found that the connectivity was enhanced from the lPFC toward the FFA during switch relative to repeat trials of the face task (i.e. switch-to-face versus repeat-face trials; Fig. 2D *left*), and also enhanced from the lPFC toward the PPA in switch relative to repeat trials of the place task (Fig. 2D *right*). These results are in line with the well-known role of the left lPFC in behavioral flexibility (Konishi et al. 2002; Crone et al. 2006; Derrfuss et al. 2005; Yeung et al. 20026; Kim et al. 2012; Jimura and Braver 2010; Tsumura et al. 2021), and suggest top-down signaling from the prefrontal cortex to stimulus-modality-dependent occipitotemporal regions during task switching (Tsumura et al. 2021).

We then asked a critical question whether the top-down signaling from the lPFC to the stimulus-modality-dependent regions is modulated depending on the uncertainty of the relevant task (i.e., multi-task level) that was manipulated by the motion coherence of task cue. During face tasks with high-coherence cue, the task-related effective connectivity was enhanced from the MT and FFA regions to the lPFC, and the directionality of connectivity between these regions was reversed in low-coherence trials (Fig. 2E *left*). Likewise, the effective connectivity was enhanced from the MT and PPA regions to the lPFC during the place task with the high coherent cue, and the directionality of the connectivity was also reversed in low-coherence trials (Fig. 2E *right*).

In order to test the reliability and robustness of the results above, we estimated the effective connectivity by 1) using an alternative estimation method (Figs. S1A-B), 2) changing the number of the regions of interest (ROIs) in the models (Figs. S1C-D), and 3) changing the definition of the ROIs (Figs. S1E-F) (see Materials and Methods). Overall results were maintained, confirming that the connectivity results were robust against estimation procedures, model structures, and model parameters. Task-unrelated intrinsic connectivity is shown in Figs S1G-H.

These collective results of effective connectivity suggest that, when relevant task was indicated ambiguously, top-down signal from prefrontal regions become stronger toward stimulus-modality-dependent occipitotemporal regions (i.e. MT and FFA/PPA for face/place tasks). On the other hand, when task cue information is more evident, bottom-up signals from the occipitotemporal regions to the prefrontal regions become stronger.

### Whole-brain decoding by a CNN classifier

Given the brain regions associated with tasks, switching tasks, and perceptual decision-making identified by univariate activation analysis and their effective connectivity mechanisms, we explored brain regions that code relevant task information using a convolutional neural network (CNN) classifier (LeCun et al. 2015). More specifically, we examined whether brain activity patterns involve discriminable information about face and place tasks during task switching with cue uncertainty, and then identified brain regions involving critical information about the relevant task.

The current analysis used VGG16 (Simonyan et al. 2015) that was trained to classify a concrete object image dataset provided by ImageNet (http://www.image-net.org/; Krizhevsky et al. 2012; Fig. S2A). We re-trained the VGG16-ImageNet model using flat whole cortical activation maps (Figs. 3A and S2B) such that it classified face and place tasks based on fine-tuning (Donahue et al. 2014; Fig. S2C). The retraining was performed based on cortical maps that were independent of the tested maps. We retrained the model using flat maps during a working memory (WM) task for face and place stimuli obtained from the Human Connectome Project (HCP) (Fig. S2C *top*; see also Materials and Methods), followed by additional retraining based on flat activation maps during target-only trials of the current task in which only the face/place target stimulus was presented without the dot cue stimulus (Figs. 1D and S2C *bottom*).

Classification accuracy for target-only trials was 82.1 ± 5.0% (mean ± SD with 10-fold cross validation), which was significantly greater than chance level (P < .001) (Fig S3A; see Materials and Methods). Interestingly, direct retraining of VGG16-ImageNet model to classify the current target-only trial maps showed little increase in accuracy (Fig. S3A). Training of a randomly initialized VGG model to classify HCP WM maps was also not successful (Fig. S3B). Thus, these results demonstrate that the current two-step retraining of the VGG16-ImageNet model was sufficient for the CNN model to learn from small sample data sets of brain images.

Given that the CNN model successfully classified face and place tasks during the target only trials with high accuracy, we then examined the classification accuracy for cue trials (Fig. S2D). Accuracy was higher than chance level in switch trials at all coherence levels [80% switch: t(28) = 4.7, P < .001; 40% switch: t(28) = 5.5, P < .001; 20% switch: t(28) = 5.4, P < .001] and 80%-repeat trials [t(28) = 3.6, P < .005] (Fig. 3B), which ensured those maps contained information about performed tasks. More importantly, classification accuracy was higher in switch than in repeat trials [F(1, 28) = 10.9; P <. 005], suggesting that cortical activation patterns involves more information about task dimension in switch than in repeat trials, although the coherence effect was absent [F(1, 28) = 0.3; P = 0.6].

We explored brain regions involving critical information to classify face and place tasks by visualizing weights of convolution layers of the CNN model. We used Grad-CAM (Selvaraju et al. 2017) that aggregates weights across convolution layers, and highlights image locations with greater weights when important information to classify the image are involved (Fig. 3C). The weight maps were created for each of the tested images (i.e. flat activation maps for cue trials), and then collected for each of the tasks (face/place), switching conditions (switch/repeat), and coherence levels (high/mid/low) (see Materials and Methods).

We first contrasted weight maps between switch and repeat trials, and calculated pixel-wise group-level z-statistics with participants treated as a random effect (Fig. 3D). Prefrontal, parietal and occipitotemporal areas including FFA and PPA (see Figs. 2C and S4 for references) showed greater weights in switch trials than repeat trials (Fig. 3D). In particular, weights became greater in trials switching to face task but not to place task in the FFA (Fig. 3D *left*). On the other hand, greater weights were observed in trials switching to the place task but not to face task in the PPA (Fig. 3D *right*). These results suggest that modality-dependent FFA and PPA encode task-relevant information to a greater degree during the switch to the task that demands stimulus discrimination of optimal modality. Next we examined the coherence effect during cue trials by calculating the weighted sum of weights for each pixel (Fig. 3E). The FFA and PPA also showed greater weights in high-coherence trials than low-coherence trials in both of the face and place tasks.

In order to statistically test dissociated weight patterns in the FFA and PPA during cue trials, ROI analysis was performed (Fig. 3F). ROIs were defined based on target only trials, independently of cue trials (see Materials and Methods). In the FFA, weights were greater during the face task than the place task [F(1, 28) = 15.2, P < 01], and in the PPA, weights were greater during the place task than the face task [F(1, 28) = 66.5, P < .001]. For weight magnitudes with optimal task-region relation (i.e., face task in FFA and place task in PPA), weights became greater in switch relative to repeat trials [F(1, 28) = 6.4, P < .05], and in high relative to low-coherence trials [F(1, 28) = 13.4, P < .01 with linear contrast]; however, there was no without switching-by-coherence interaction [F(1, 28) = 0.1, P = .75]. In a separate analysis, we used a leave-one-subject-out procedure when retraining the classifier to classify the activation maps of the target only trials, and then tested the remaining subject (Fig. S5). Overall results are consistent, suggesting that subject-specific noises are not dominant in our results.

These results suggest that the FFA and PPA involve more modality specialized task-related pattern information in high-coherence trials. Thus, modality-dependent occipitotemporal regions may encode relevant task information (i.e. FFA for face task and PPA for place task), which is enhanced in switch trials with a high-coherence task cue.

### Whole-brain decoding by SVM

In order to complement decoding and mapping by the CNN classifier, we performed another decoding analysis using an SVM classifier. Similar to the CNN classifier analysis above, the classifier was trained based on HCP WM task such that the classifier discriminates the dimension of the tasks (face or place) (see Materials and Methods). We then tested the experimental data to examine classification performance for face and place tasks of the cue trials.

We found that accuracy was higher than chance level in switch trials at all coherence levels [80% switch: t(28) = 4.5, P < .001; 40% switch: t(28) = 2.5, P < .05; 20% switch: t(28) = 3.3, P < .005; Fig. 4A], and higher in switch than repeat trials [F(1, 28) = 5.0; P <. 05], which is consistent with the CNN classifier results.

Weights of the SVM classifier were mapped onto 2D cortical surface of the brain in order to identify brain regions with greater weights to classify the face and place tasks. Occipitotemporal regions including the FFA and PPA showed prominent reverse directed weights (Fig. 4B), indicating that these regions involve important information to classify the two tasks. Notably, these maps are consistent with the CNN-based mapping, especially in the FFA and PPA (Figs. 3D/E). We also trained another SVM based on target-only trials in the current task, and found that the classification accuracy of cue trials (Fig. S6A) and weight maps (Fig. S6B) were consistent to those with the CNN classifier (Figs. 3B/D-E) and whole-brain SVM (Figs. 4A-B). We note that SVM weight maps (Figs. 4B and S6B) reflect a hyperplane calculated by training data (i.e. maps for HCP WM or current target-only trials), whereas CNN weight maps reflect degree of contribution to classify tested image (i.e. current cue trial maps) (see Discussion).

### Decoding mapping by searchlight SVM

The above CNN and SVM classifiers were based on pattern information of whole brain cortical regions. Another SVM analysis was also performed using the searchlight procedure (see Materials and Methods). By exploring across the whole brain, searchlight was used to identify brain regions where local image voxels involved pattern information about the performed task to classify face and place tasks. Again, the classifier in each searchlight was trained using HCP datasets, and thus the training and testing datasets were independent.

For target-only trials, classification accuracy was significantly higher in the FFA and PPA regions (Fig. 4C and Table S4), suggesting that modality-dependent occipitotemporal regions involve relevant task information; this result is consistent with the univariate analysis (Fig. 2C).

Voxel-wise classification accuracy maps were contrasted between switch vs. repeat trials for each participant, and group-level statistical tests were performed in order to identify brain regions where discriminable pattern information is greater in switch trials than in repeat trials. Occipitotemporal regions showed a significant effect of switching (Fig. 4D and Table S5), indicating that, in these regions, classification accuracy is higher in switch relative to repeat trials, consistent with our previous study Tsumura et al. 2021). Interestingly, these regions were spatially located in-between the FFA and PPA, where pattern information of searchlight classifiers may modestly involve both of face-related and place-related information, possibly in a balanced manner (Tsumura et al. 2021). These regions also showed a coherence effect with higher accuracy in high-coherence trials (Fig. 4E and Table S6). Notably, these results were consistent with weight mapping of the CNN classifier (Figs. 3D-E).

It is important that these regions showed significantly higher classification accuracy than chance-level in switch and repeat trials (Figs. 7A-B and Tables S7-8), and cue trials at each coherence level (Figs. S7C-E and Tables S9-11). This assures that the differential accuracies between switch and repeat trials, and across coherence levels were attributable to accuracy enhancement in switch trials with the high-coherence cue.

These collective results suggest that occipitotemporal regions adjacent to stimulus-modality-dependent FFA/PPA areas involves information about ongoing task, and that the information amount is increased during task switching with more coherent cue presentation. These results are also consistent with those of the classification performance and mapping based on whole-brain CNN and SVM classifiers. Such differential classification accuracy was not observed in fronto-parietal regions well known to be involved in executive control (Figs. S7F-G and Tables S5-6), even when the classifier was trained by target-only trials in the current experiment (Figs. S7H-I).

## Discussion

The current study examined neural mechanisms during task switching under situation where task cue involved uncertainty. Task-related neural coding in FFA/PPA became more evident during task switching and also when the relevant task was cued more explicitly. When task-cue was distinct, the lPFC received task-related signals from the MT region and PPA/FFA, and the direction of the signal was reversed when the task cue involved more ambiguity. These results suggest a distributed cortical network of fronto-occipitotemporal regions for behavioral flexibility where task-related signal among these regions help to implement task representation depending on the ambiguities of external cue (Fig. 5).

**Figure 5.**
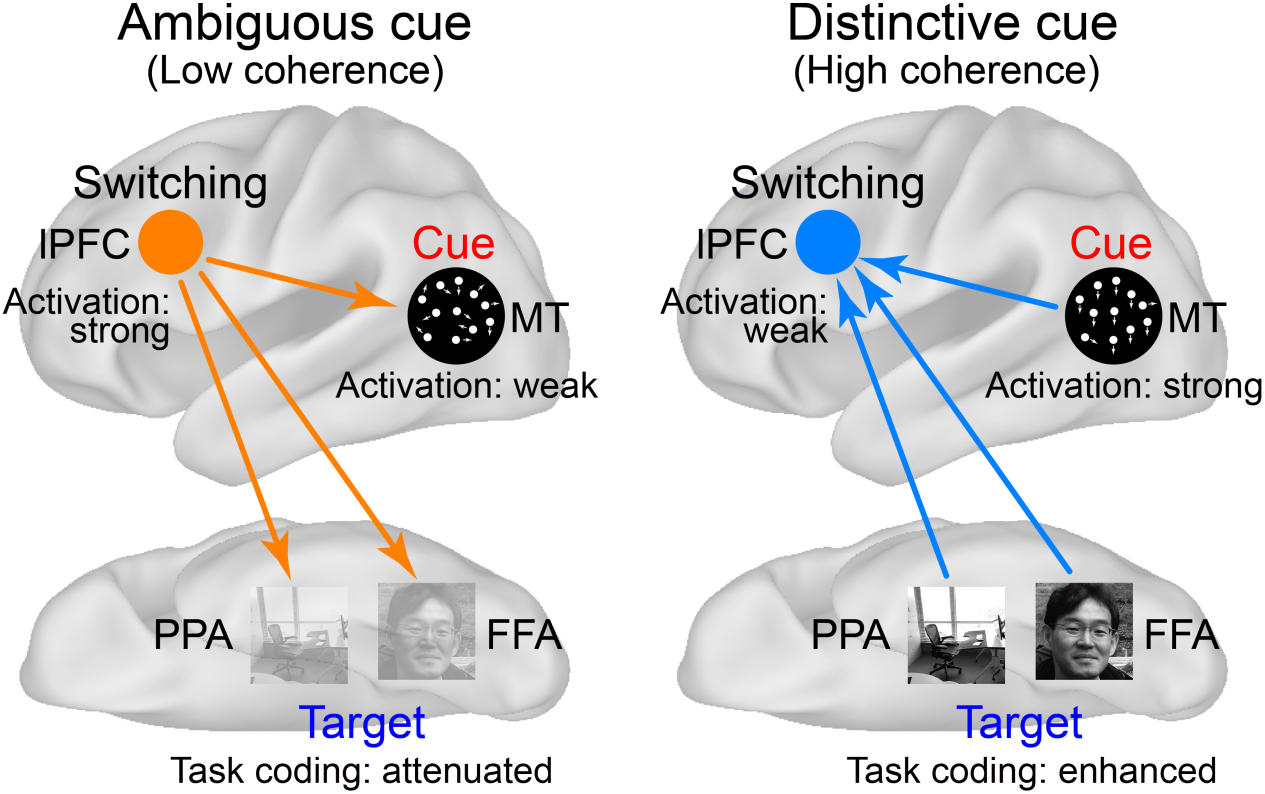
Putative model of behavioral flexibility under perceptual uncertainty. Schematic diagrams for functional mechanism among the middle temporal (MT) region, fusiform face area (FFA), parahippocampal place area (PPA), and lateral prefrontal cortex (lPFC) during task switching with cue uncertainty. The arrows indicate signal directions.

### Neural mechanism for task switching and perceptual decision making

Prior work of task switching has examined switch-related neural mechanisms under situations where perceptual uncertainties were applied to target stimulus (Kayser et al. 2010; Mante et al. 2013; Zhang et al. 2013; Kumano et al. 2016; Tsumura et al. 2021). In contrast, the current study manipulated uncertainty of the task cue that involves multi-task information (Figs. 1C/D and 5). Notably, the task cue indicates relevant task dimension at the upper layers of the hierarchical task set structure (Koechlin et al. 2003; Bunge et al. 2005; Badre 2008; Jimura and Braver 2010; Fig 1B), and thus the current study allowed us to elucidate higher-level cognitive functions governing task switching and perceptual decision-making.

For putative mechanisms to achieve task switching under cue uncertainty, three hypotheses are possible: 1) a unitary mechanism implements task switching under perceptual uncertainty in a task cue; 2) distinct mechanisms for perceptual decision-making and task switching interactively guides successful task switching; and 3) a hub-like region links the two distinct mechanisms. Behaviorally, an interaction between switching and coherence levels was absent (Figs. 1E/F; see also Results); this is consistent with the univariate imaging analysis and CNN classifier mapping showing no interaction effect. The absence of an interaction effect, together with distinct brain regions associated with cue/target perception and task switching (Figs. 2A-C and 3D-F), suggests distributed mechanisms for task switching under cue ambiguities, which supports the second hypothesis. Thus, by modulating effective connectivity and interregional signaling depending on cue ambiguity, these regions cooperatively guide behavioral flexibility (Fig. 5). This interpretation is also compatible with the central role of frontal regions for flexible task control (Egner and Hirsch, 2005; Kayser et al. 2010; Cole et al. 2013; Waskom et al. 2014).

The current results and interpretations are consistent with our recent study (Tsumura et al. 2021) in that complementary cortical mechanisms are engaged in task switching under situations where goal-relevant information is ambiguous, but extend it by showing homologous and more elaborated mechanisms help to achieve task switching when perceptual uncertainty is involved in the task cue instead of the target stimuli. The complexity and sophistication of the mechanisms is attributable to the increased number of occipitotemporal regions: in the current study, three regions, MT, FFA, and PPA, are involved, whereas two regions, MT and ventral visual complex (VVC), are involved in our prior study (Tsumura et al. 2021).

### Fronto-occipitotemporal network mechanisms

DCM analyses revealed top-down signal from the lPFC to the FFA or PPA, depending on the task to be switched, and this top-down signaling is enhanced when task cue involved ambiguity during switching (Fig. 5). The top-down mechanisms may reflect supplemental attention to visual stimulus required to collect task cue information about the tasks to be performed (Lee and D’Esposito 2012; Zanto et al. 2011). The supplemental attention involving frontal engagement may complement stimulus-modality-dependent activation in occipitotemporal regions (Desimone and Duncan 1995; Kastner and Ungerleider 2000; Corbetta and Shulman 2002; Lewis-Peacock and Postle 2008).

In contrast, bottom-up signaling with increased cue information may reflect the conversion of sensory information to behavioral information through an information stream from the visual sensory area to executive control areas (Desimone and Duncan 1995; Kastner and Ungerleider 2000). Thus, when the task cue was apparent, the bottom-up signal was strengthened because cue-related information is more available, which may help to enhance task switching performance.

On the other hand, such signal reversal did not occur in our prior study, and the top-down signals from prefrontal to occipitotemporal regions complement switching independently of the uncertainty of the target stimulus (Tsumura et al. 2021). It is possible that the signal reversal is characteristic of the uncertainty of the cue stimulus, as the cue stimulus indicates more abstract information in the upper layers of the task-set hierarchy (Fig. 1B). Thus, the uncertainty of the abstract information may implicate more complex mechanisms to implement task sets to be performed (Koechlin et al. 2003; Bunge et al. 2005; Badre 2008; Jimura and Braver 2010).

### Comparisons of classification and mapping among machine learning techniques

In the current study, whole brain exploration of task-related neural representation was performed by three approaches based on three machine learning techniques, 1) CNN classifier, 2) whole-brain cortical SVM, and 3) searchlight SVM.

CNN classified activation maps along task dimensions based on all pixels across whole cortical regions, and the classification accuracy was higher in switch trials than repeat trials. One novel signature of the current CNN classifier approach is that brain regions involving critical information to classify cue trials were mapped by aggregated weight gradients across convolution layers based on the Grad-CAM technique. It is notable that this CNN weight mapping is available on image-by-image basis for testing data, which is not the case for the SVM mapping using whole cortical images. Then, the CNN weight mapping revealed that task representation in the FFA during the face task and in the PPA during the place task was enhanced in switch and more coherent trials than in repeat and less coherent trials. Increased task-related activation (Figs. 2C and S4) may be associated with higher classification accuracy and enhanced task representation.

Standard voxel-wise univariate GLM analysis identifies brain regions where the MRI signal is differentiated between task conditions, but does not necessarily indicate that identified brain regions are critical for task performance; this makes it hard to identify brain regions playing an important role in cognitive functions (i.e., reverse inference). In contrast, the current CNN classifier demonstrated that the visualization of convolution layers of the classifier for brain activation is useful in identifying brain regions that characterize task performance.

The analysis based on the CNN classifier was complemented by a standard SVM analysis for whole brain cortical maps that tested identical map images. The classification accuracy of cue trials was highly consistent between the two classifiers. Additionally, SVM weight maps also showed differential weights in the FFA and PPA, which is also consistent with the CNN classifier. One notable technical limitation of the SVM is that weight mapping for testing classification for cue trials was unavailable, unlike Grad-CAM of CNN classifier. Thus the SVM weight map indicates that the FFA and PPA is critical to classify tasks for the HCP N-back working memory task or target only trials in the current task (i.e. training data), but not necessarily for cue trials of the current task (i.e. testing data). Nonetheless, together with differential classification accuracy among cue conditions, the SVM weight maps suggest that pattern information in the FFA and PPA is distinct (i.e. distant from separating hyperplane) in switch trials than in repeat trials.

Another approach to identify brain regions that characterize task performance is the searchlight SVM, which also allows whole-brain exploration of activation patterns, but individual classifications were restricted in local brain regions (Figs 4C-E and S7A-E). This is in contrast to the CNN classifier and whole-brain cortical SVM that are trained and classify based on a whole brain image. Nonetheless, results were complementary to those whole-brain-based classifiers in that 1) occipitotemporal regions adjacent to the FFA and PPA were capable of task classification, and 2) the classification performance in these regions became higher in switch and more coherent trials.

The higher classification accuracy in switch trial is consistent with our recent study (Tsumura et al. 2021) whereas previous MVPA studies of task switching suggested that task coding in fronto-parietal regions is attenuated in a switch trial (Qiao et al., 2017), and task coding is independent of task switching (Loose et al. 2017).

Interestingly, those previous studies used task-cueing paradigms, where a task cue was presented in each trial (Loose et al. 2017; Qiao et al. 2017); conversely, the current study and our recent study used an intermittent cue paradigm in which the switch trial occurred after successive correct trials for the alternative task without presenting a cue (Fig 1C/D; Tsumura et al. 2021). The variability in the cueing procedures among the studies may yield the variability of classification accuracy.

### Classifier training using independent open resource data

One notable analysis procedure in the current machine-learning-based functional brain mapping is that classifier was trained using an open resource dataset that was independently collected from the current experiment. This procedure ensured independence between the training and testing data.

Task switch and working memory may involve distinct cognitive control demands, with the former related to behavioral flexibility (Allport et al. 1994; Rogers and Monsell 1995) and the latter related to active maintenance and updates of goal-relevant information (D’Esposito and Postle 2015). However, the two tasks used common visual stimulus categories (face and place); thus, perceptual demands may involve some degree of commonality. Recognition demands for the presented stimuli were also distinct: the current task-switching paradigm used male-female and indoor-outdoor discriminations during face and place tasks, respectively, but HCP working memory tasks used discrimination of identicalness to past stimuli. Additionally, the current task used face-place superimposed stimuli, and thus the identical stimulus set was used during face and place tasks, whereas HCP working memory task used distinct visual stimulus sets during face and place blocks. Thus task representation examined in the current study may reflect visual perception or attention rather than low level visual features.

For the CNN classifier, training involved two steps: retraining of HCP working memory maps, and additional retraining of target-only trials. CNN classifies maps based on whole cortical areas including fronto-parietal regions in which differential sub-regions are recruited during task switching (Dove et al. 2000; Rushworth et al. 2002; Bunge et al. 2005; Crone et al. 2006; Derrfuss et al. 2005; Kim et al. 2012; Yeung et al. 2006; Malagon-Vina et al. 2018; Bissonette et al. 2017; Fouragnan et al. 2019; Nee et al. 2016; Koechlin et al. 2003; Jimura and Braver 2010; Tsumura et al. 2021) and working memory (Courtney et al. 1997; Miller and Cohen 2001; D’Esposito and Postle 2015). Thus additional retraining of the CNN model based on identical recognition demands (i.e. target-only trials) was effective in classifying tasks to optimize the CNN model for classification using whole cortical images. Distinct weight differences in the parietal cortex (Figs. 3D-E) may partially be attributable to higher performance with the additional retraining.

Because incremental training of HCP working memory trials and the target only trials in the current study is irrelevant to SVM, these two datasets were separately trained for whole brain cortical SVM, and weight maps were consistent especially in occipitotemporal regions (Figs. 4B and S6B). Importantly, classification accuracy for the cue conditions were consistent in SVMs with the two training datasets, and also with the CNN classifier. The sample size was much smaller for the current target-only trials than HCP dataset, but classification performance was comparable between those two classifiers (Figs. 4A and S6A). Thus, SVM may thus not require a larger sample size like the HCP data for training, while CNN training needed incremental training even with large sets of image data.

The searchlight SVM using HCP working memory maps as training data identified occipitotemporal regions spatially closed to the FFA/PPA showing higher classification accuracy for target-only trials. Moreover, classification accuracy was higher in high-coherence switch trials. Interestingly, these classification results were absent in fronto-parietal regions, well known to be involved in executive control (Figs. S7F/G), even when the searchlight classifier was trained by the target-only trials of the current task (Figs. S7H/I). One possibility for this discrepancy is that control and recognition demands are incompatible while perceptual modality is compatible in HCP working memory trials, target only trials, and switch/repeat trials. Then the distinct control and recognition demands might be reflected in classification incompatibility in the fronto-parietal regions.

A CNN model successfully learned and decoded task-related fMRI images of HCP without flattening the images, and retraining of the classifier was also effective for small subset of the HCP data (Wang et al. 2020). However, this technique is unavailable for the current experiment using event-related design (see Materials and Methods for more details), and retraining of the VGG16/ImageNet model based on flattened 2D activation maps was powerful for the current dataset.

## Acknowledgements

We thank Drs. Akira Funahashi, Shori Nishimoto, and Teppei Matsui for scientific comments on the study and manuscript. We thank Ms. Maoko Yamanaka for administrative assistance.

## Author contributions

K.T. and K.J. designed the experiment and study. K.T., R.A., K.N. and K.J. collected the data. K.T., K.K. and K.J. analyzed the data. M.T., J.C. and Y.H. contributed to the analysis design of imaging data and to the development of machine learning classifier. K.T., M.T., J.C., K.N. and K.J. wrote the manuscript.

## Supplementary Material

### Supplementary Figures

**Fig. S1.**
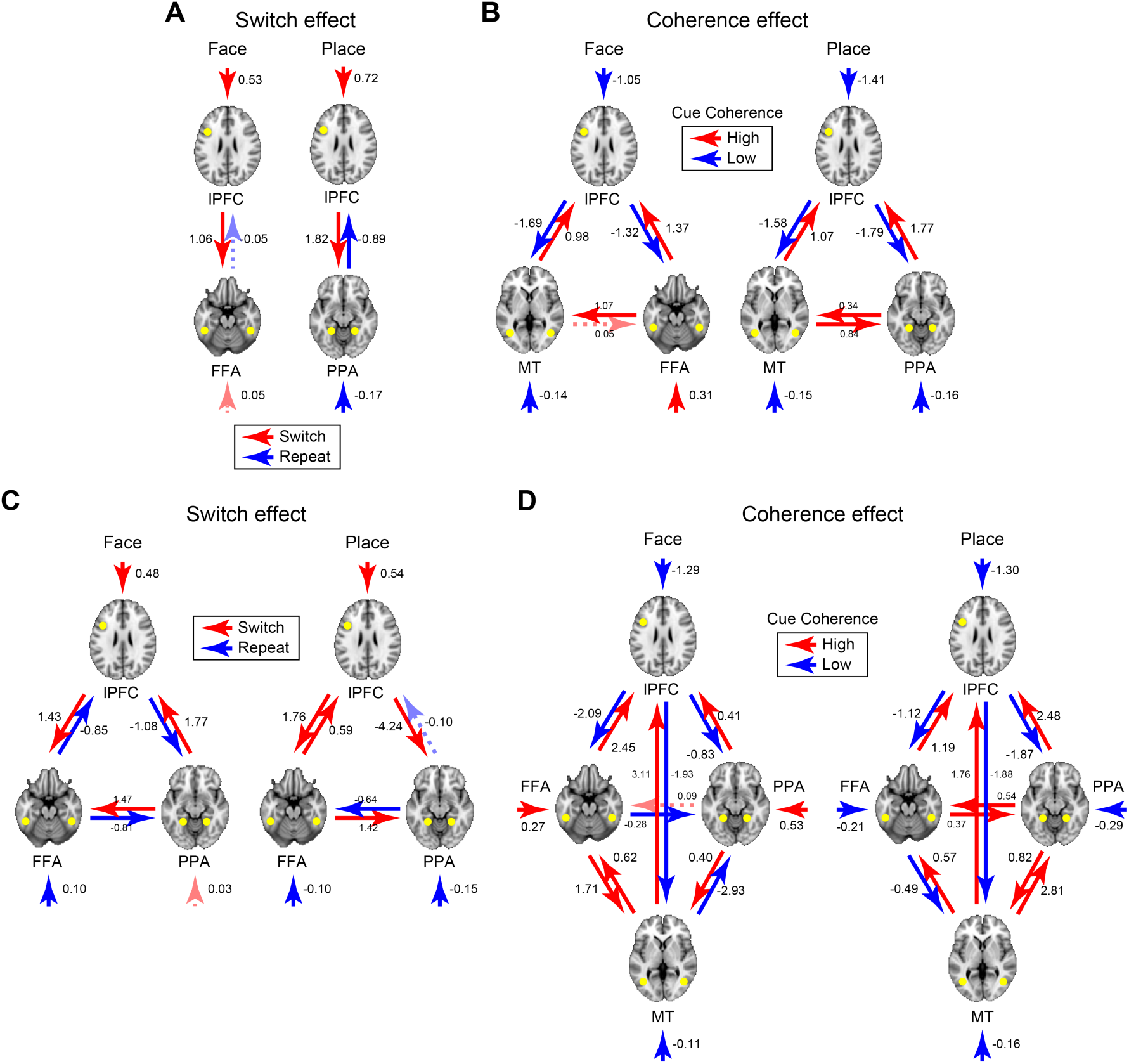
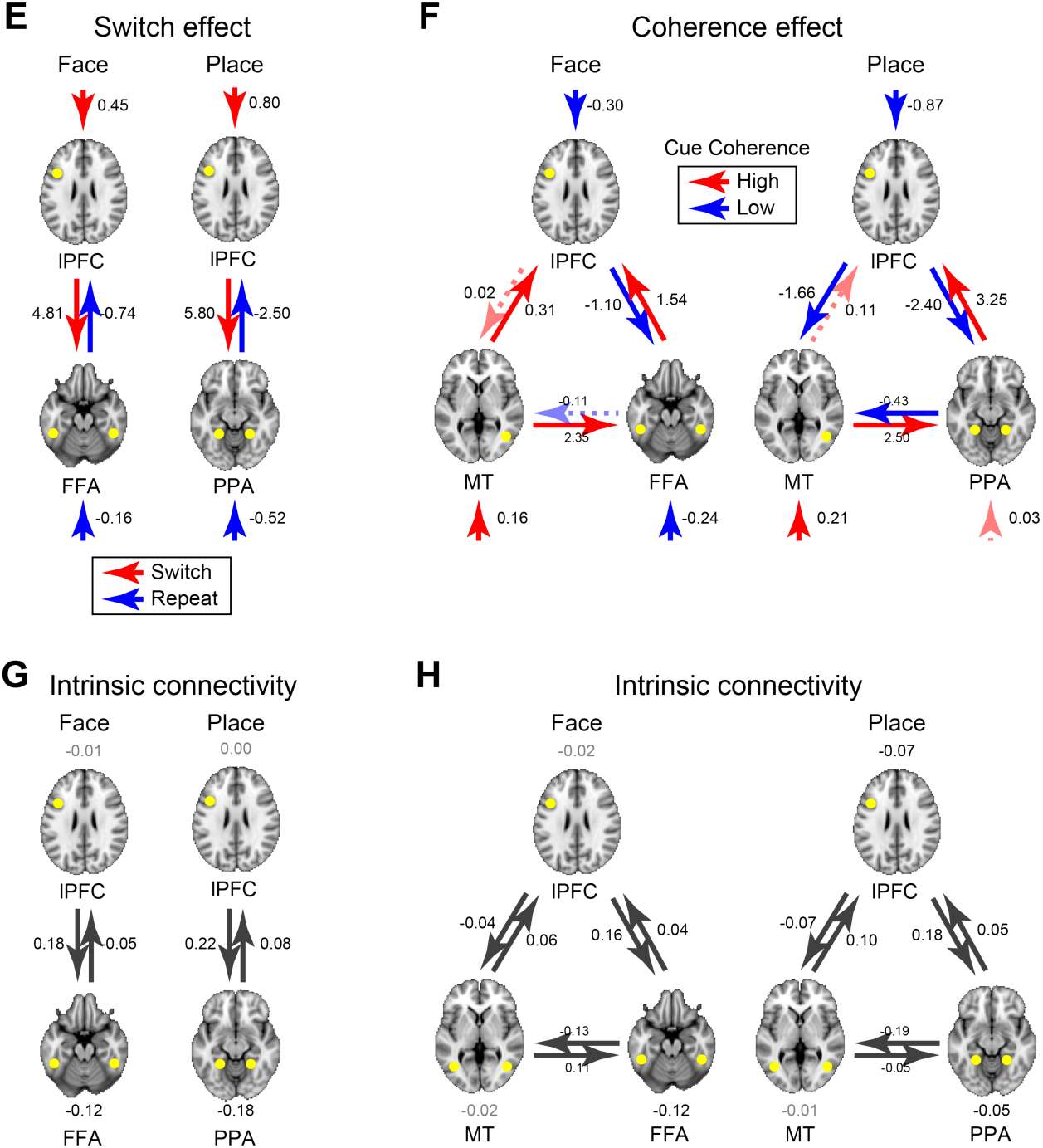
Effective connectivity analysis. (A-B) Parameters estimated without parametric empirical Bayes. (A) Switching effect. (B) Coherence effect. (C-D) The numbers of ROIs in the model was changed. (C) Switching effect was examined in a model involving the lateral prefrontal cortex (lPFC), fusiform face area (FFA), and parahippocampal place area (PPA) for both of the face and place tasks. (D) Coherence effect was examined in a model involving lPFC, FFA, PPA, and middle temporal (MT) regions. (E-F) The definition of ROIs. (E) Switching effect. (F) Coherence effect. The other analysis procedures were identical to those in Figs. 2D-E. (G-H) Task unrelated intrinsic connectivity, estimations of “A” matrix in the dynamic causal modeling. (G) Switching effect (H) coherence effect. Gray solid lines indicate statistical significance.

**Fig. S2.**
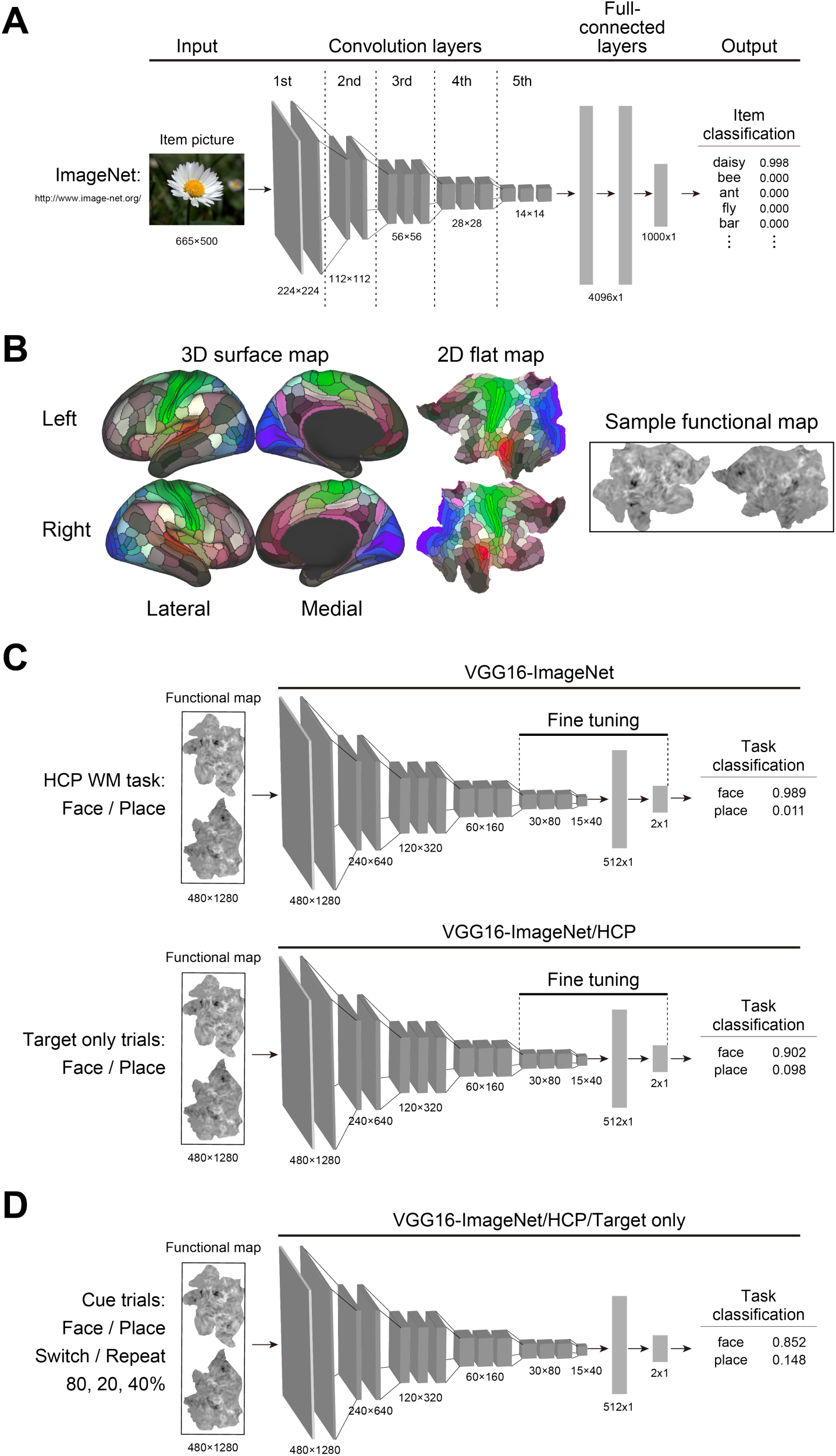
Training and testing procedure for the convolutional neural network classifier (CNN). (A) VGG16 classifier pre-trained for concrete item pictures available from ImageNet. The classifier consists of 5-tier convolution layers and 3-tier full-connected layers, and is capable of classifying pictures of concrete items into 1,000 categories. (B) 3D surface maps (*left*) are mapped to 2D flat maps (*middle*). For illustration purpose, colored areas are cortical subregions functionally and anatomically segmented in prior work (ref. 42). Sample flattened 2D z-maps for the contrast face vs. fixation and place vs. fixation (*right*). (C) Training procedures for 2D-flat images. ImageNet-pretrained VGG16 model (A) was first retrained such that it classified activation maps for the face and place task during working memory task distributed by Human Connectome Project (HCP) (top). The retrained model was further retrained such that it classifies activation maps for the face and place task during target only trials in the current experiment (*bottom*). (D) Testing procedure. The classifier trained by the two stepwise procedure (C) was tested for classification of face and place tasks during cued trials involving switch and repeat trials with different coherence levels.

**Fig. S3.**
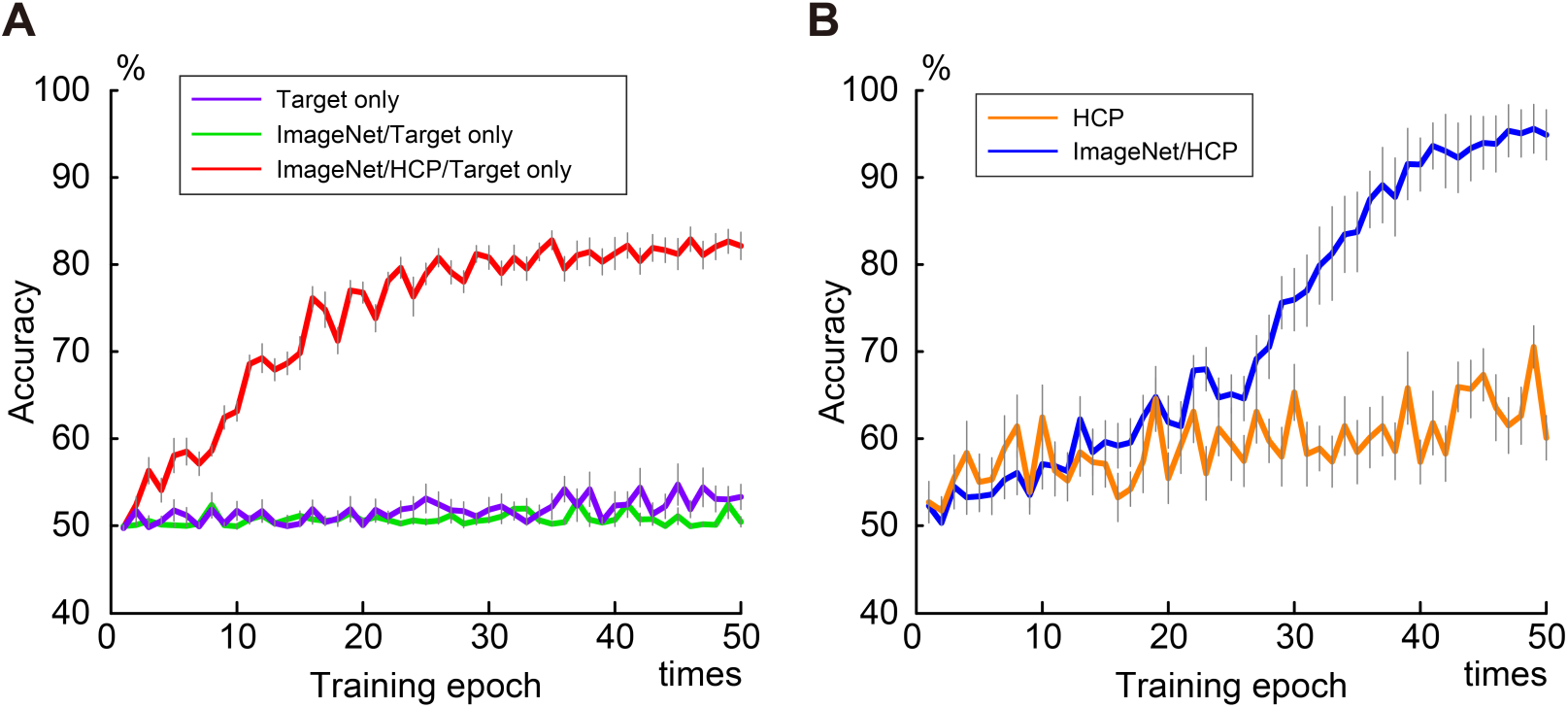
Classification accuracy of convolutional neural network during training epoch. Vertical and horizontal axes indicate validation accuracy, and training epoch, respectively. (A) Validation accuracy for face/place classification of the current target-only trials. Purple line indicates the learning curve of a model trained based on the activation maps of target-only trials from a randomly initialized parameter. Green line indicates the learning curve of a model re-trained by the activation maps of target-only trials from pre-trained parameter by ImageNet. Red line indicates the learning curve of a model re-trained by the activation maps of target-only trials from the pre-trained parameter by Human Connectome Project (HCP) working memory maps and ImageNet (i.e. green line). Error bars indicate standard error of the mean across cross-validation testing. (B) Validation accuracy for face/place classification of working memory trials provided by HCP. Orange line indicates the learning curve of a model trained based on the activation maps of working memory trials from a randomly initialized parameter. Blue line indicates the learning curve of a model trained by activation maps of HCP working memory trials from a pre-trained parameter by ImageNet. Error bars indicate standard error of the mean across cross-validation testing.

**Fig. S4.**
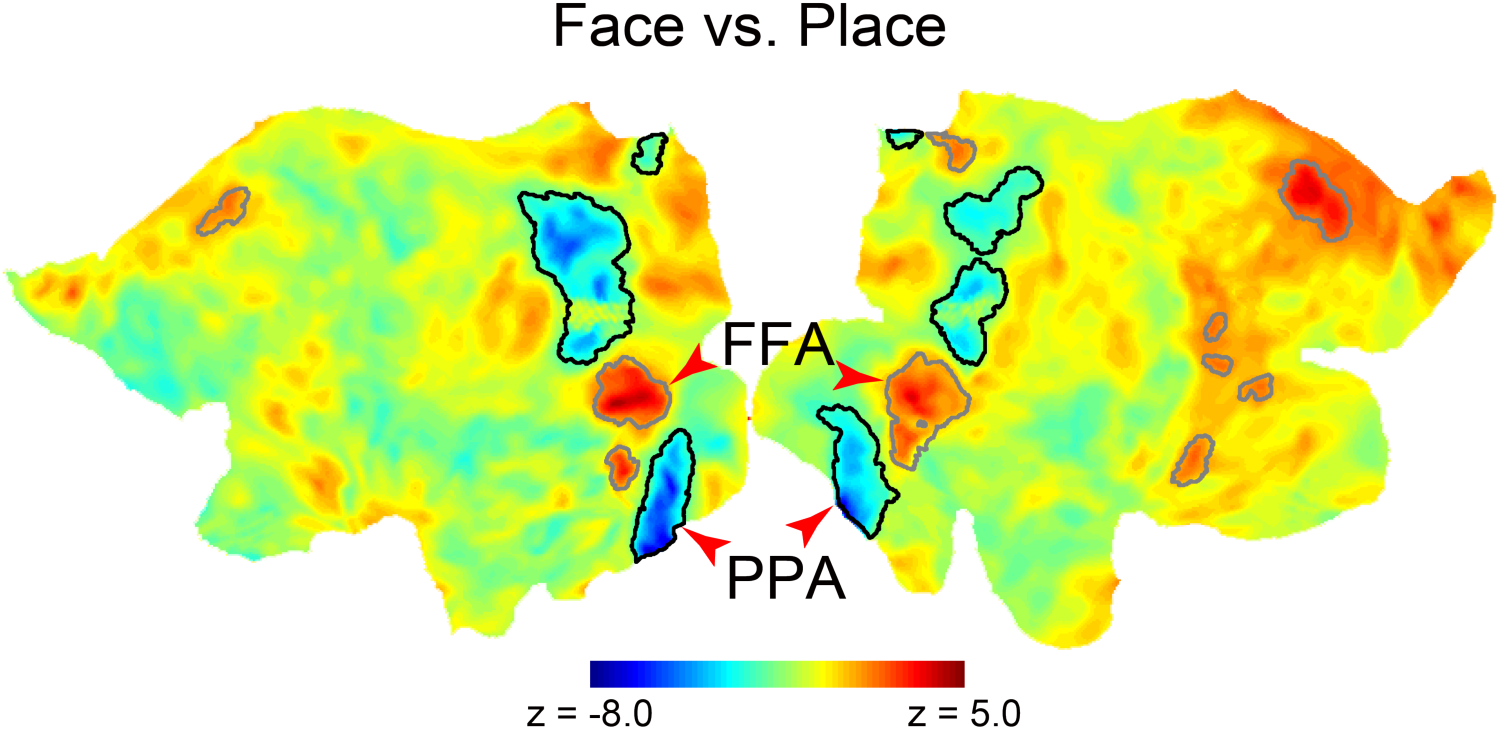
Statistical activation z-map for contrast face vs. place tasks in univariate analysis. Data are derived from those shown in Fig. 2C, but maps are overlaid onto 2D-flat map of the cortical surface of the brain. Hot and cool colors indicate greater activity in face and place tasks, respectively. Closed lines indicate prominent activity during face (gray) and place (black), respectively, as identified by a group-level analysis (Table S3). The fusiform face area (FFA) and parahippocampal place area (PPA) are indicated by red arrow heads.

**Fig. S5.**
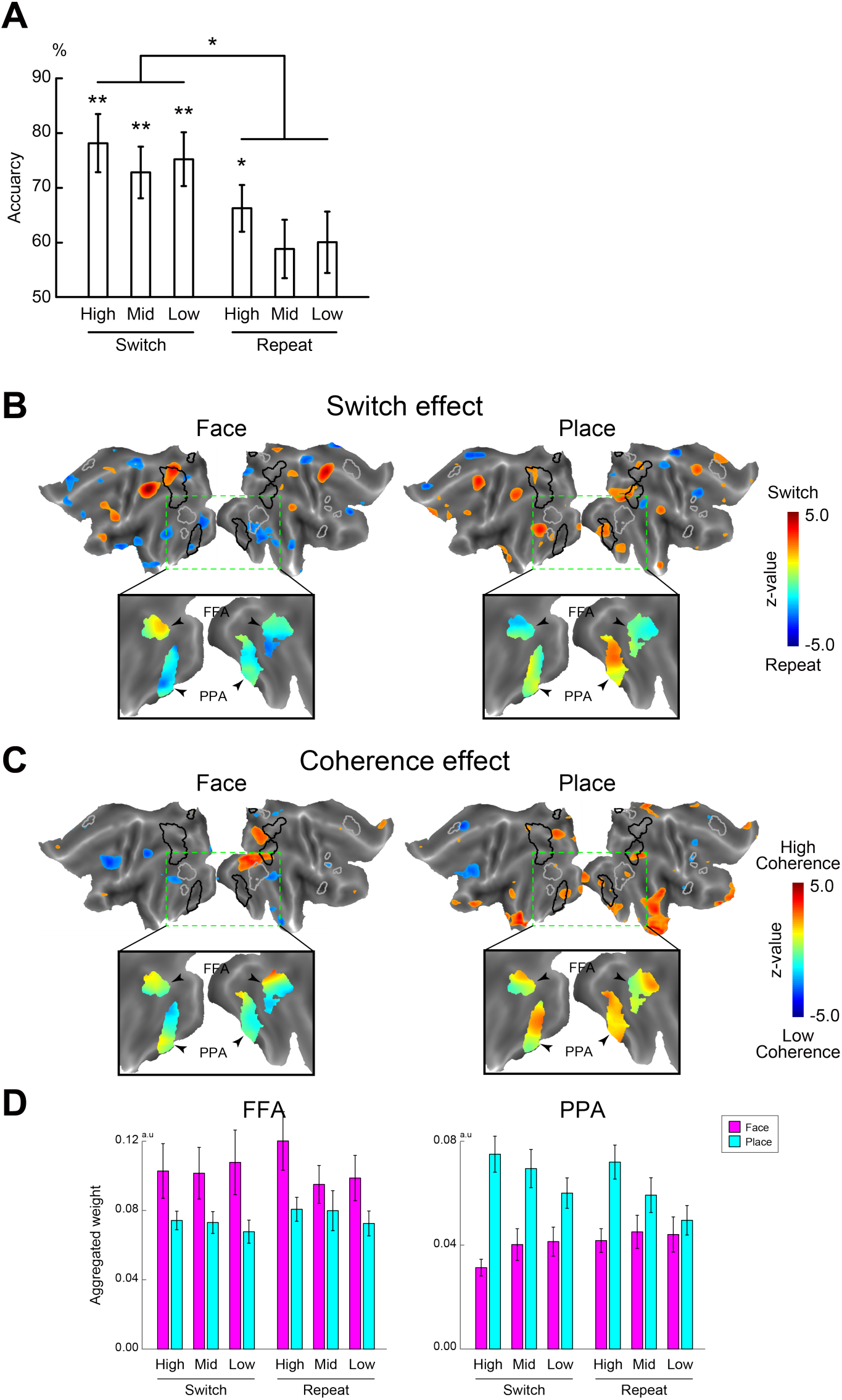
Classification accuracy and weight maps of CNN classifier. A leave one-subject-out procedure was used in training and validation of the CNN classifier. (A) Classification accuracy for each task condition. (B) Visualization of aggregated weight contrasts for switch vs. repeat trials in face task (*left*) and place task (*right*). Occipitotemporal regions in rectangular boxes with green broken lines were expanded below. (C) Visualization of weight contrast for coherence effect in face task (*left*) and place task (*right*). (D) Regions of interests (ROIs) analysis. All formats are similar to those in Figs. 3B/D-F.

**Fig. S6.**
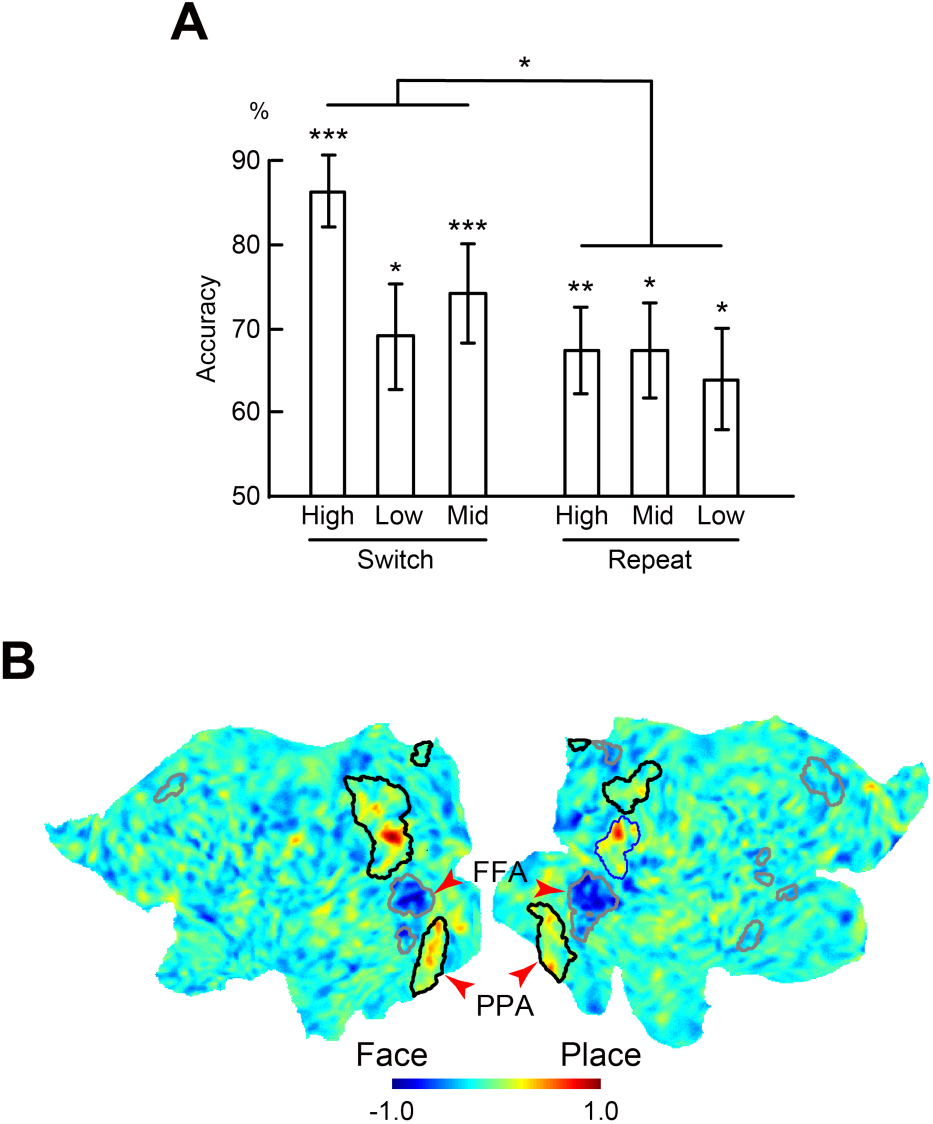
The support vector machine classifier trained by target-only trials. (A) Classification accuracy for cue trials. Error bars indicate standard error of the mean across participants. Formats are similar to those in Fig. 2B. (B) Weight map of SVM. Maps are overlaid onto flat maps of the cortical brain. Hot and cool colors indicate high and low weights, respectively. Gray and black lines overlaid on flat map indicate clusters significantly activated during face and place tasks in univariate analysis, respectively (Figs. 2C and S4). FFA and PPA was indicated by red arrow heads.

**Fig. S7.**
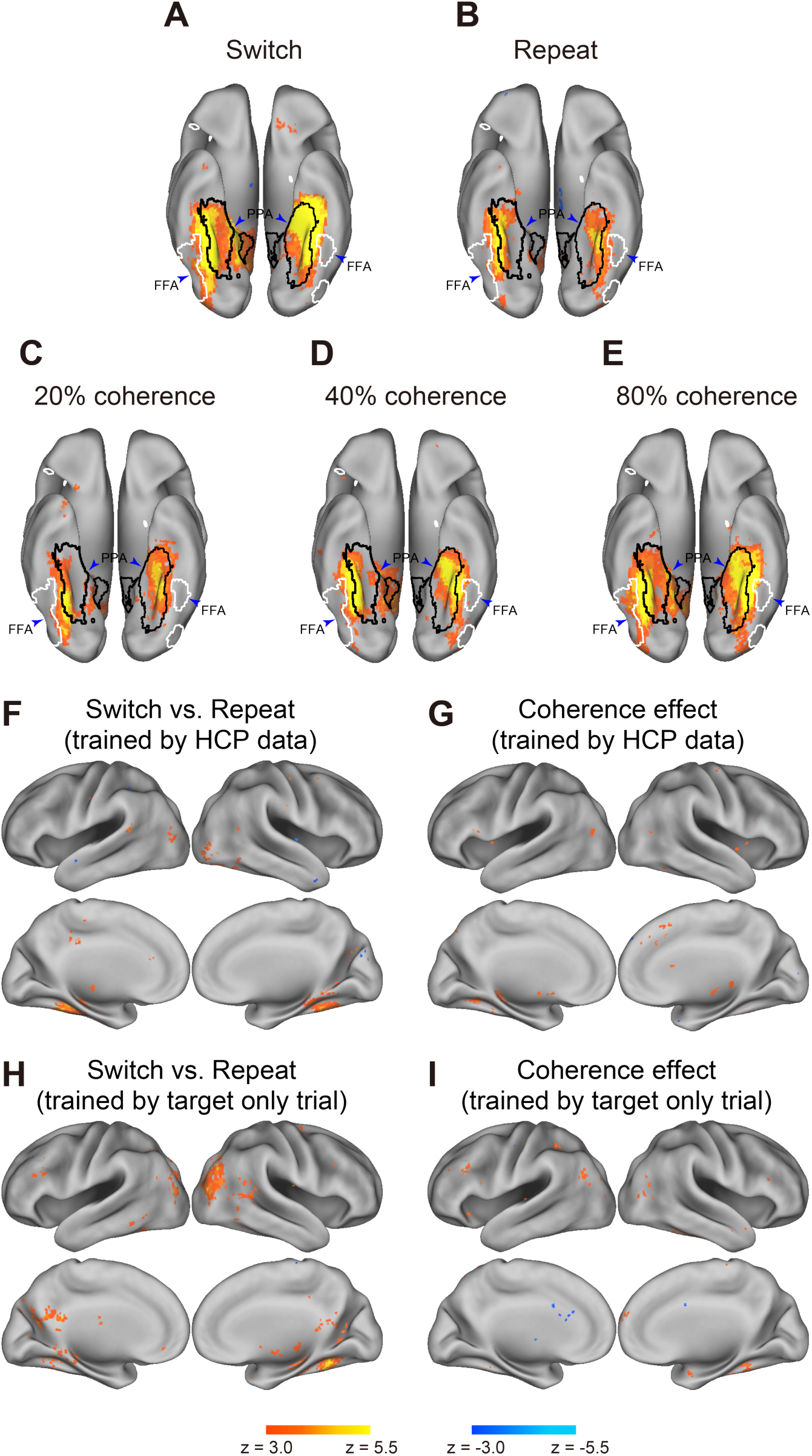
Searchlight multi-variate pattern analysis. Hot and cool colors indicate statistical level for classification accuracy relative to chance level. Maps are overlaid onto 3D surface of the brain and displayed from a ventral view. White and black closed lines overlaid onto 3D surface of the brain indicate significant cluster for contrast face vs. place tasks in univariate analysis, respectively (Figs. 2C and S4). The fusiform face area (FFA) and parahippocampal place area (PPA) are indicated by blue arrow heads. (A) Switch trial. (B) Repeat trial. (C) 20% coherence trial. (D) 40% coherence trial. (E) 80% coherence trial. (F) Accuracy difference between switch and repeat trials with a model trained by Human Connectome Project (HCP) data. Hot and cool colors indicate higher accuracy in switch and repeat trials, respectively. (G) Differential classification accuracy depending on coherence effect with a model trained by HCP data. Hot and cool colors indicate higher accuracy in high and low coherent trials, respectively. (H) Accuracy difference between switch and repeat trials with model trained by target-only trials of the current dataset. (I) Differential classification accuracy depending on the coherence effect with model trained data of target-only trials. The formats are similar to those in panel (F).

### Supplementary Tables

**Table S1.**
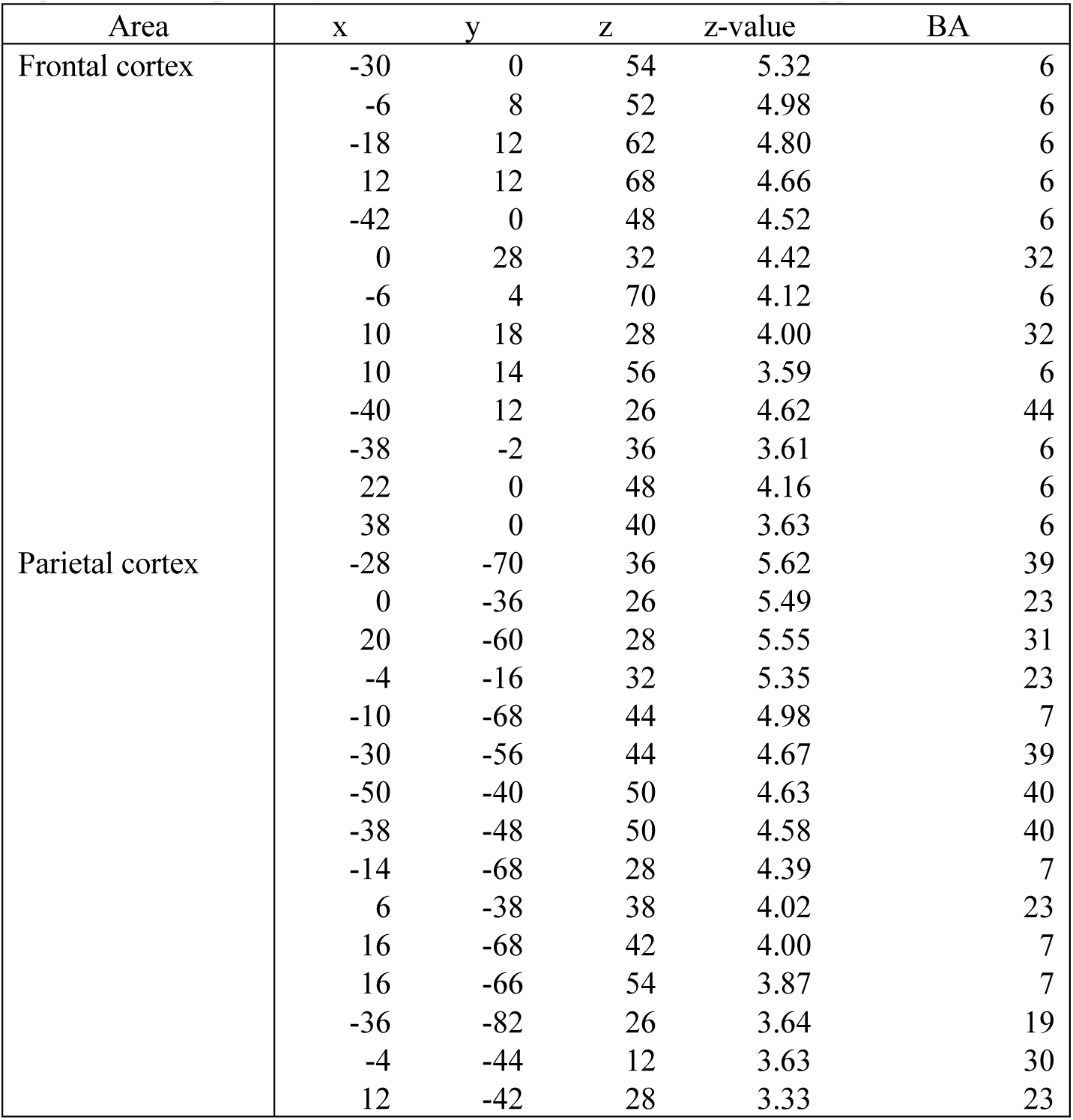
Brain regions showing significant signal increase and decrease in the contrast of switch vs. repeat trials. Coordinates are listed in MNI space. Positive and negative z-values indicate increase in switch and repeat trials, respectively. BA indicates Brodmann areas and is approximate.

**Table S2.** Brain regions showing significant parametrical effect with motion coherence. Coordinates are listed in MNI space. Positive and negative z-values indicate increase in high and low coherent trials, respectively. *: corrected within motion-related brain regions (see Methods).

| Area | x | y | z | z-value | BA |
| --- | --- | --- | --- | --- | --- |
| Frontal cortex | 0 | 18 | 50 | -5.90 | 8 |
|  | 10 | 24 | 46 | -5.59 | 8 |
|  | 54 | 20 | 38 | -5.43 | 9/44 |
|  | 46 | 14 | 4 | -5.43 | 44 |
|  | 52 | 14 | 16 | -5.21 | 44 |
|  | -56 | 12 | 28 | -5.16 | 44 |
|  | 36 | 4 | 54 | -4.62 | 6 |
|  | 48 | 6 | 34 | -4.62 | 6 |
|  | -32 | 0 | 50 | -4.49 | 6 |
|  | -38 | 10 | 32 | -4.43 | 8 |
|  | 8 | 30 | 28 | -4.41 | 8 |
|  | 42 | 24 | 28 | -4.39 | 9 |
|  | 2 | 42 | -16 | 4.37 | 11 |
|  | 4 | 30 | -18 | 4.33 | 11 |
|  | 20 | 12 | 46 | -4.31 | 6 |
|  | -50 | 8 | 10 | -4.25 | 44 |
|  | -42 | 24 | 26 | -4.08 | 9/44 |
|  | 38 | 12 | 20 | -3.82 | 44 |
|  | 40 | 36 | -4 | -3.82 | 47 |
|  | -8 | 46 | -6 | 3.80 | 10 |
|  | 24 | 8 | 62 | -3.79 | 6 |
|  | 34 | 34 | 34 | -3.78 | 9 |
|  | -52 | 18 | -6 | -3.67 | 47 |
|  | 48 | 38 | 14 | -3.35 | 46 |
|  | -44 | 38 | 28 | -3.21 | 9/10 |
| Parietal cortex | -18 | -56 | 74 | 4.70 | 7 |
|  | -16 | -48 | 36 | 4.37 | 31 |
|  | -12 | -68 | 30 | -4.28 | 7 |
|  | -24 | -64 | 38 | -4.25 | 39 |
|  | -30 | -52 | 46 | -4.19 | 7/39 |
|  | 14 | -48 | 18 | -4.19 | 23 |
|  | 58 | -46 | 36 | -4.08 | 39/40 |
|  | -48 | -40 | 48 | -4.07 | 40 |
|  | 26 | -40 | 64 | 4.03 | 1/5 |
|  | 20 | -58 | 26 | -4.02 | 23 |
|  | 42 | -50 | 52 | -3.86 | 40 |
|  | -30 | -42 | 56 | 3.85 | 5/7 |
|  | -40 | -46 | 40 | -3.66 | 39 |
|  | 6 | -46 | 32 | 3.63 | 23 |
|  | 42 | -46 | 38 | -3.50 | 39 |
|  | 32 | -42 | 46 | -3.27 | 7 |
| Temporal cortex | -42 | -54 | -8 | -5.45 | 37 |
|  | -30 | -58 | -10 | -4.10 | 37 |
|  | 56 | -64 | 2 | 3.75* | 37/19 |
| Others | 32 | 24 | 4 | -5.66 | insula |
|  | -30 | 26 | 0 | -5.56 | insula |
|  | 0 | -32 | -2 | -4.66 | brain stem |
|  | 12 | -18 | -8 | -4.30 | brain stem |
|  | 8 | -10 | 4 | -3.83 | thalamus |
|  | -16 | -28 | -4 | -3.60 | thalamus |

**Table S3.**
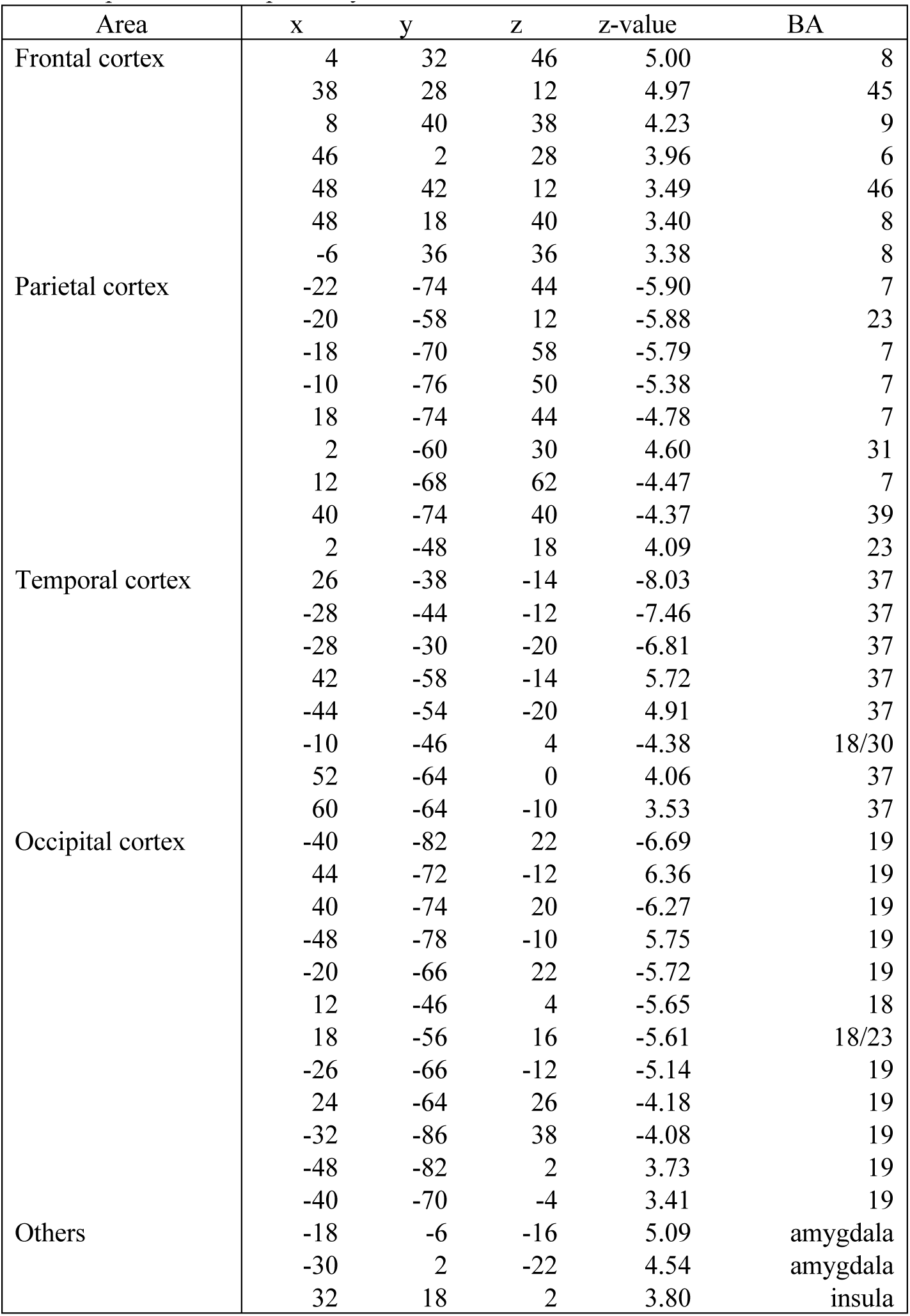
Brain regions showing significant signal increase and decrease in the contrast of face vs. place in target-only trials. Coordinates are listed in MNI space. Positive and negative z-values indicate increase in face and place trials, respectively.

**Table S4.**
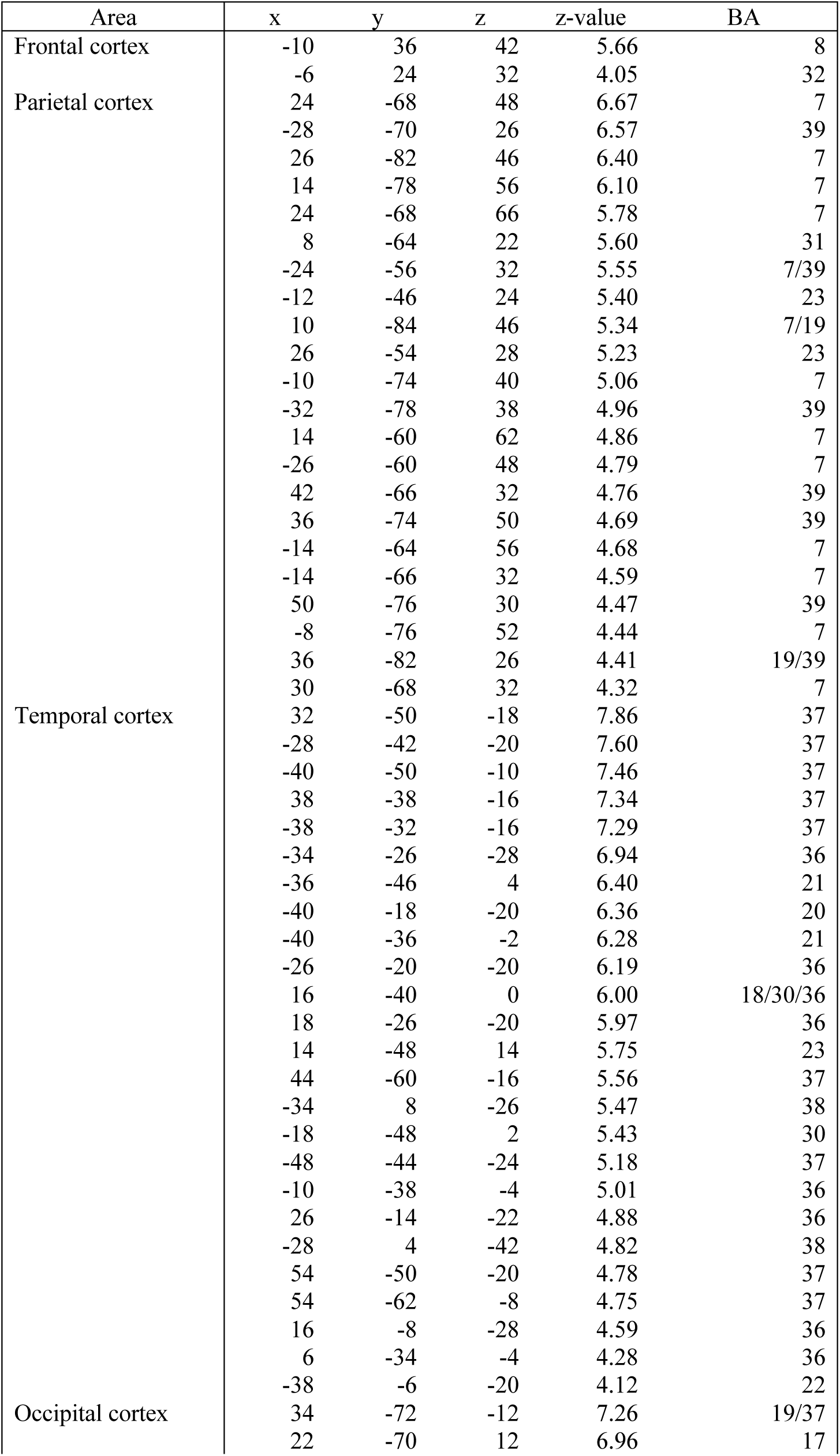
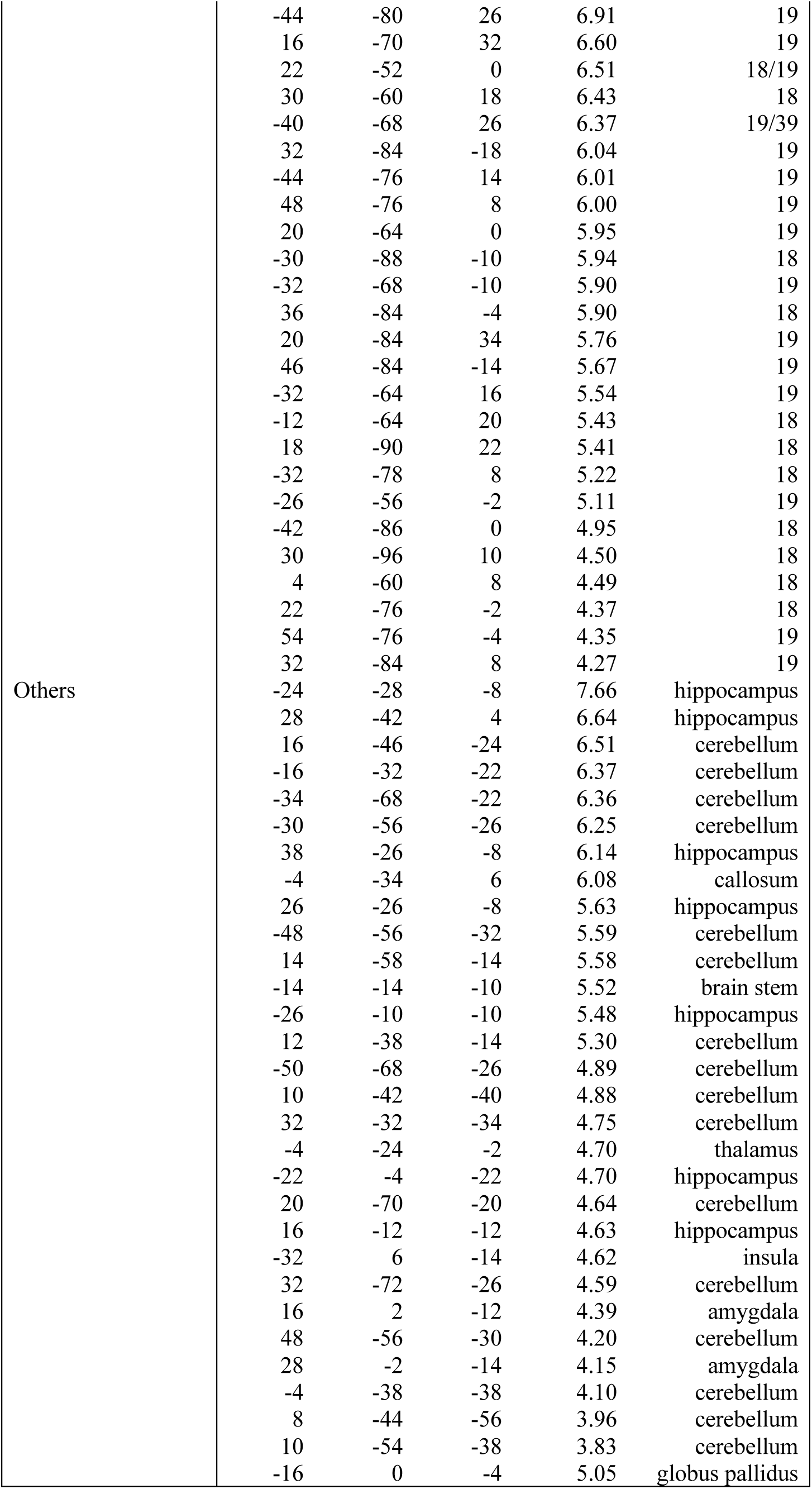
Brain regions showing significant accuracy in classifying task dimension (face/place) for target-only trials. Coordinates are listed in MNI space. Positive z-values indicate higher accuracy than chance level.

**Table S5.**
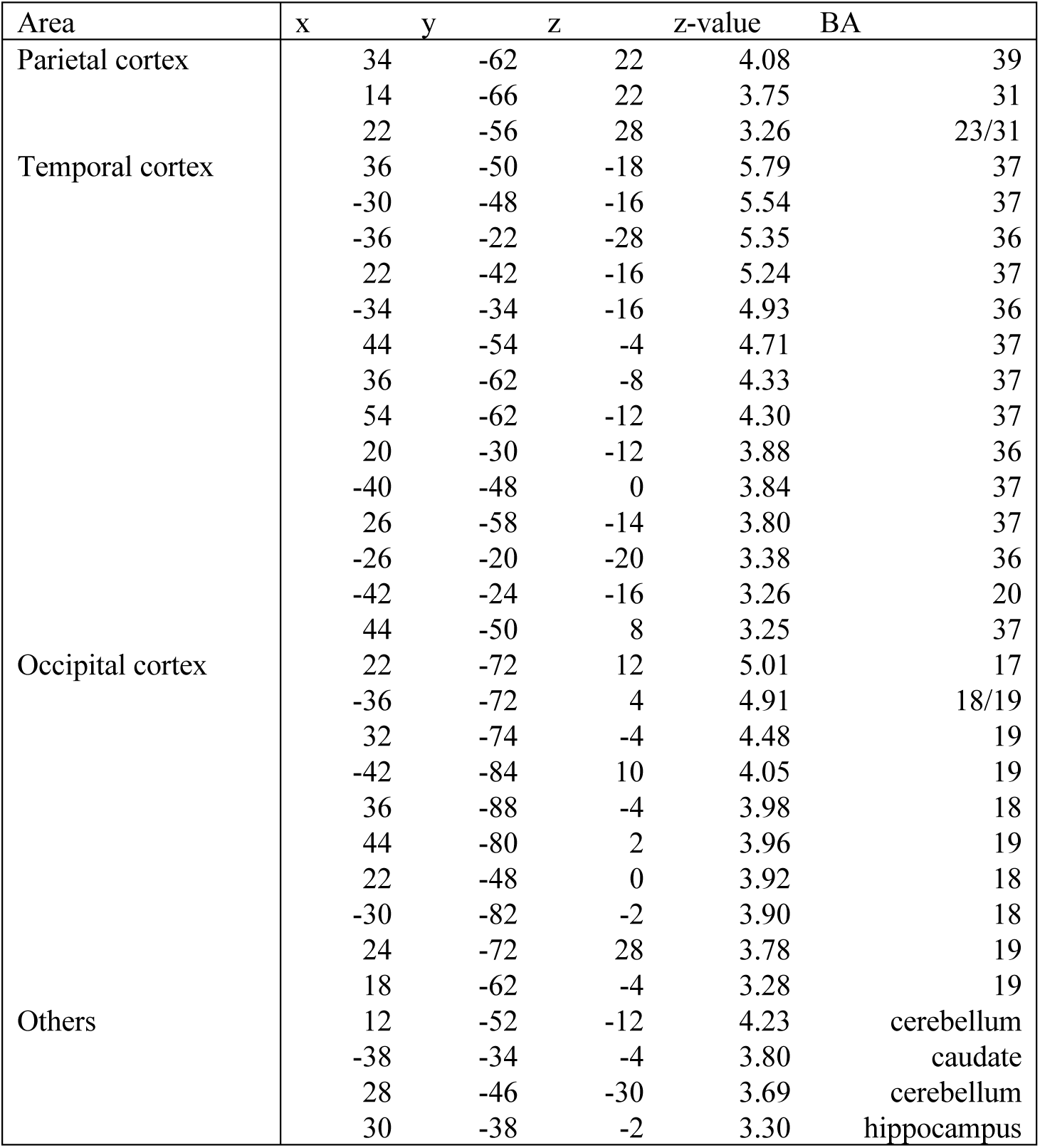
Brain regions showing significant accuracy difference for face-place classification between switch and repeat trials. Coordinates are listed in MNI space. Positive and negative z-values indicate higher accuracy in switch and repeat trials, respectively.

**Table S6.** Brain regions showing significant accuracy modulation for face and place tasks depending on coherence level. Coordinates are listed in MNI space. Positive and negative z-values indicate higher accuracy in high and low coherent trials, respectively.

| Area | x | y | z | z-value | BA |
| --- | --- | --- | --- | --- | --- |
| Temporal cortex | -30 | -62 | -8 | 4.85 | 37 |
|  | -24 | -38 | -8 | 4.29 | 36 |
|  | -34 | -52 | -20 | 4.25 | 37 |
|  | -42 | -40 | -18 | 4.20 | 37 |
|  | -14 | -28 | -12 | 3.78 | 36 |
| Occipital cortex | -18 | -54 | -8 | 3.78 | 19 |
| Others | -38 | -32 | -4 | 4.18 | caudate |
|  | -6 | -30 | 0 | 3.37 | thalamus |
|  | -34 | -46 | -32 | 3.17 | cerebellum |

**Table S7.**
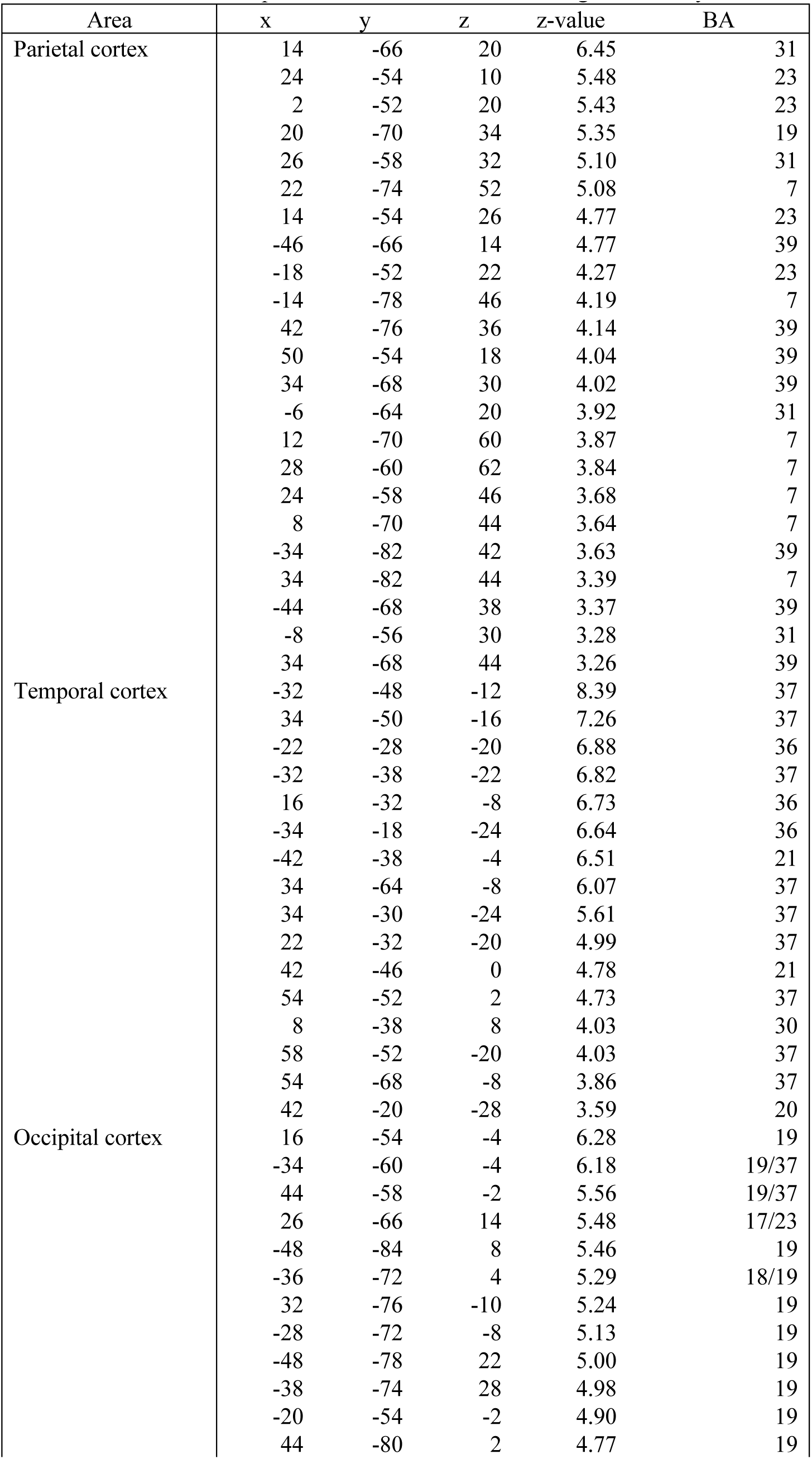
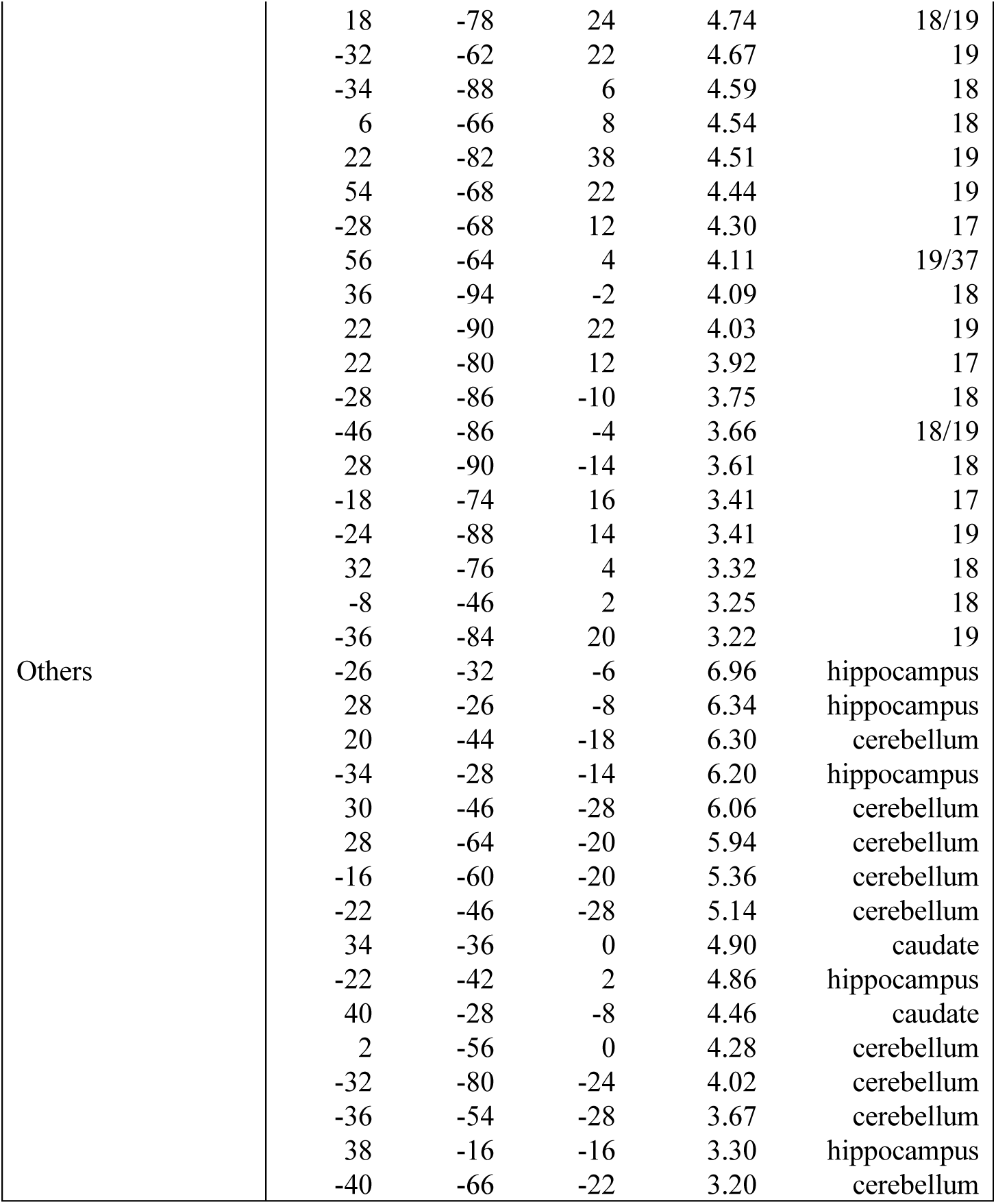
Brain regions showing significant accuracy to classify task dimension (face/place) in switch trials. Coordinates are listed in MNI space. Positive z-values indicate higher accuracy than chance level.

**Table S8.**
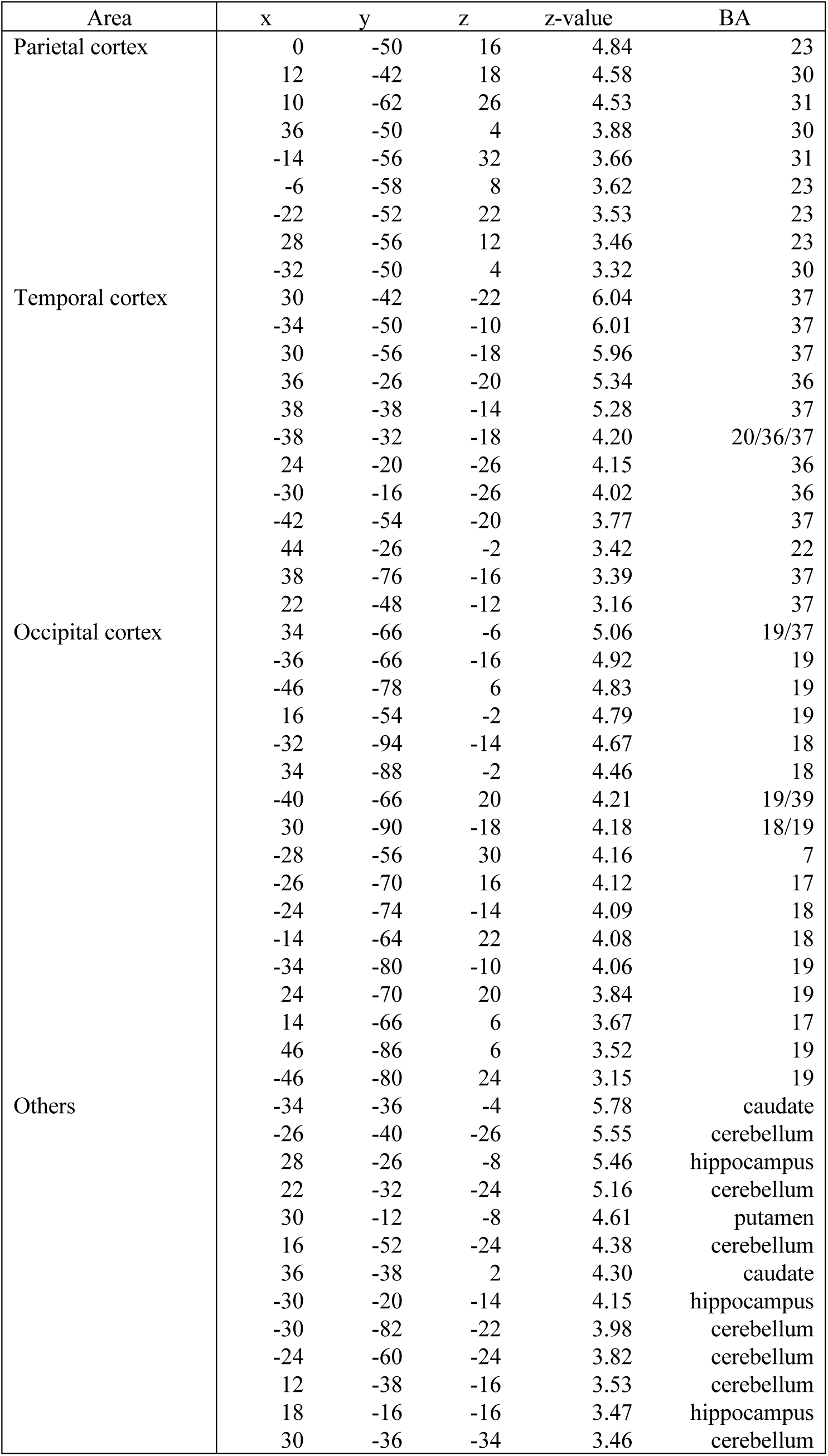
Brain regions showing significant accuracy to classify task dimension (face/place) in repeat trials. Positive z-values indicate higher accuracy than chance level.

**Table S9.**
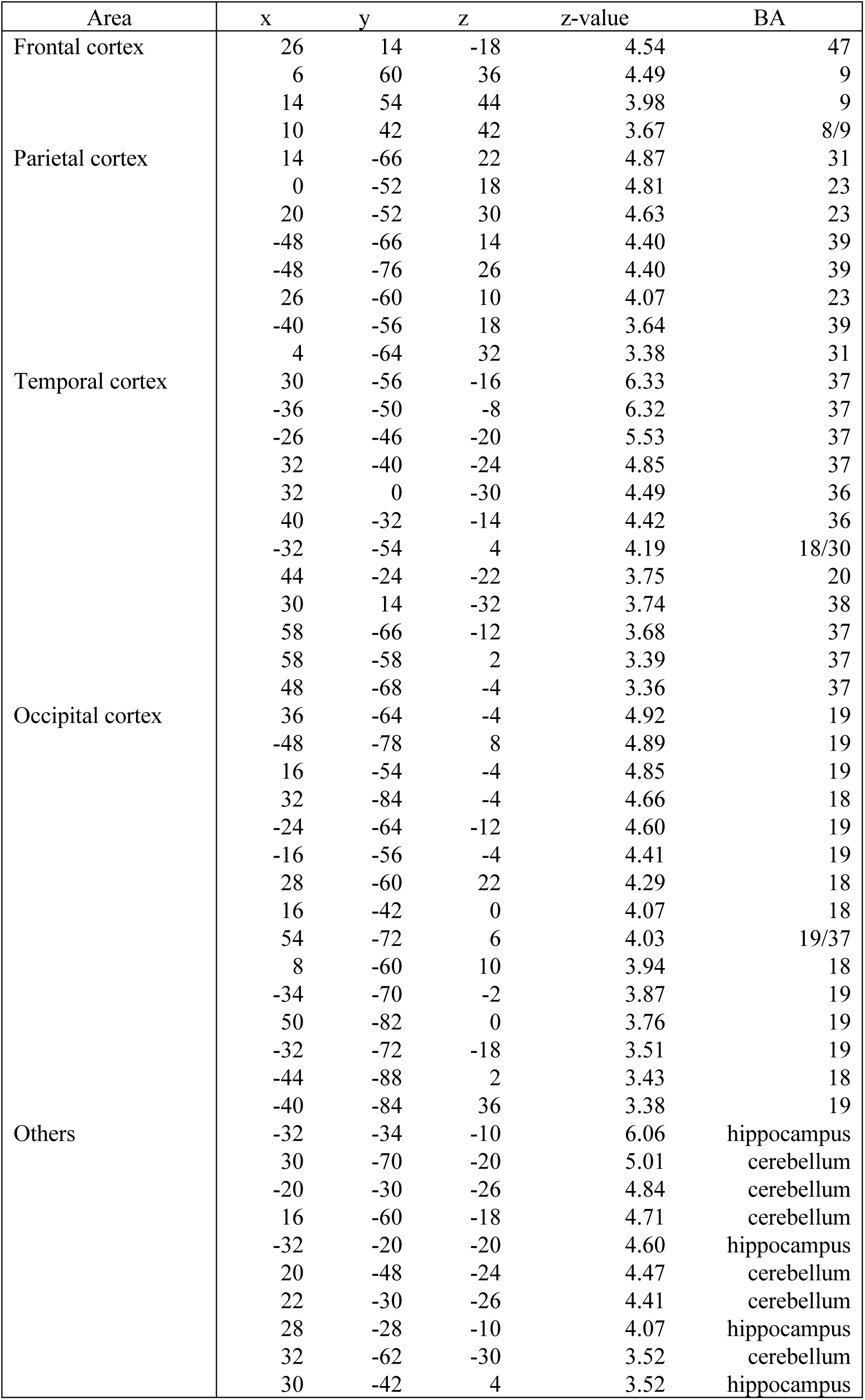
Brain regions showing significant accuracy to classify task dimension (face/place) in 20% coherence trials. Coordinates are listed in MNI space. Positive z-values indicate higher accuracy than chance level.

**Table S10.**
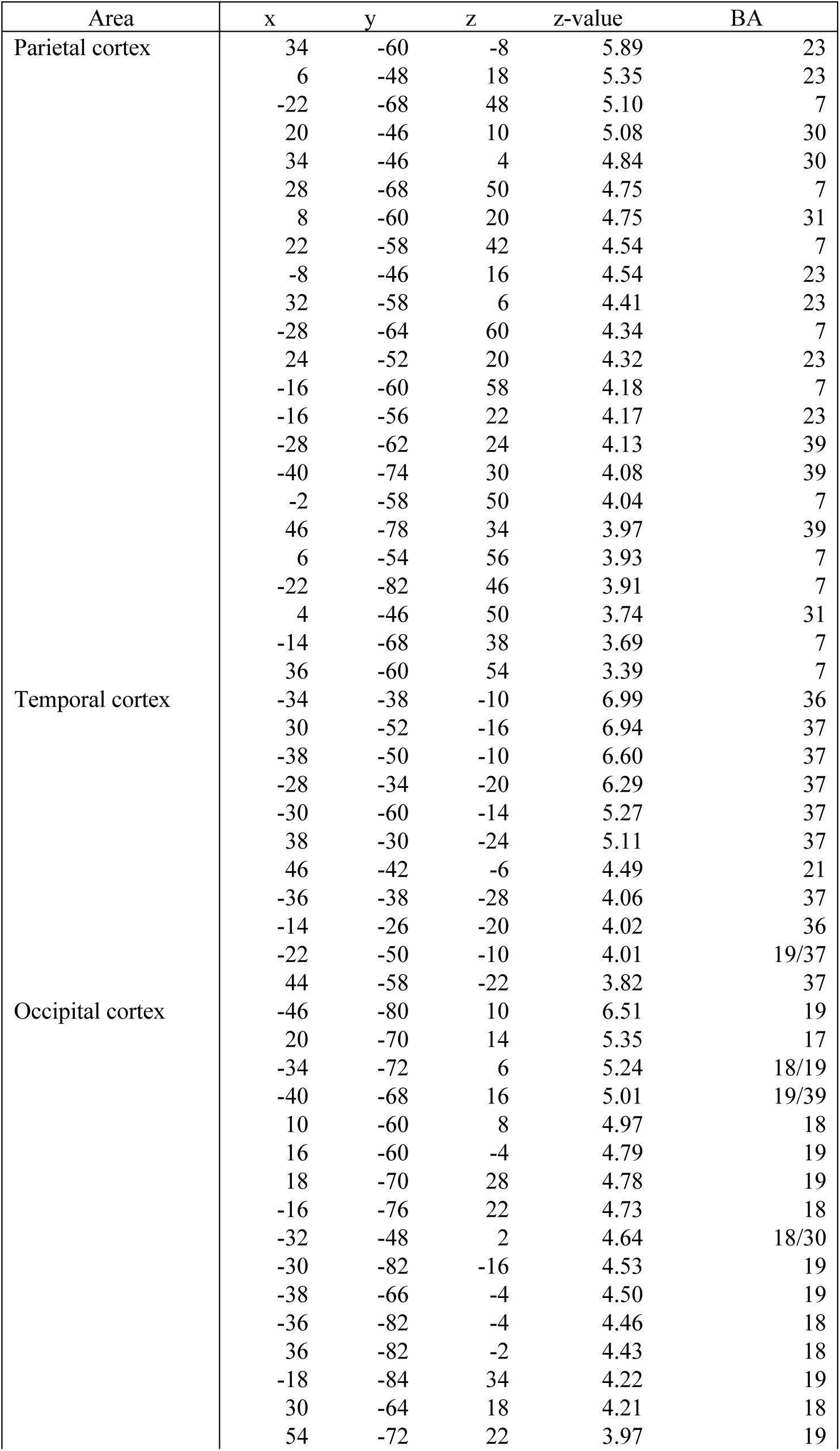
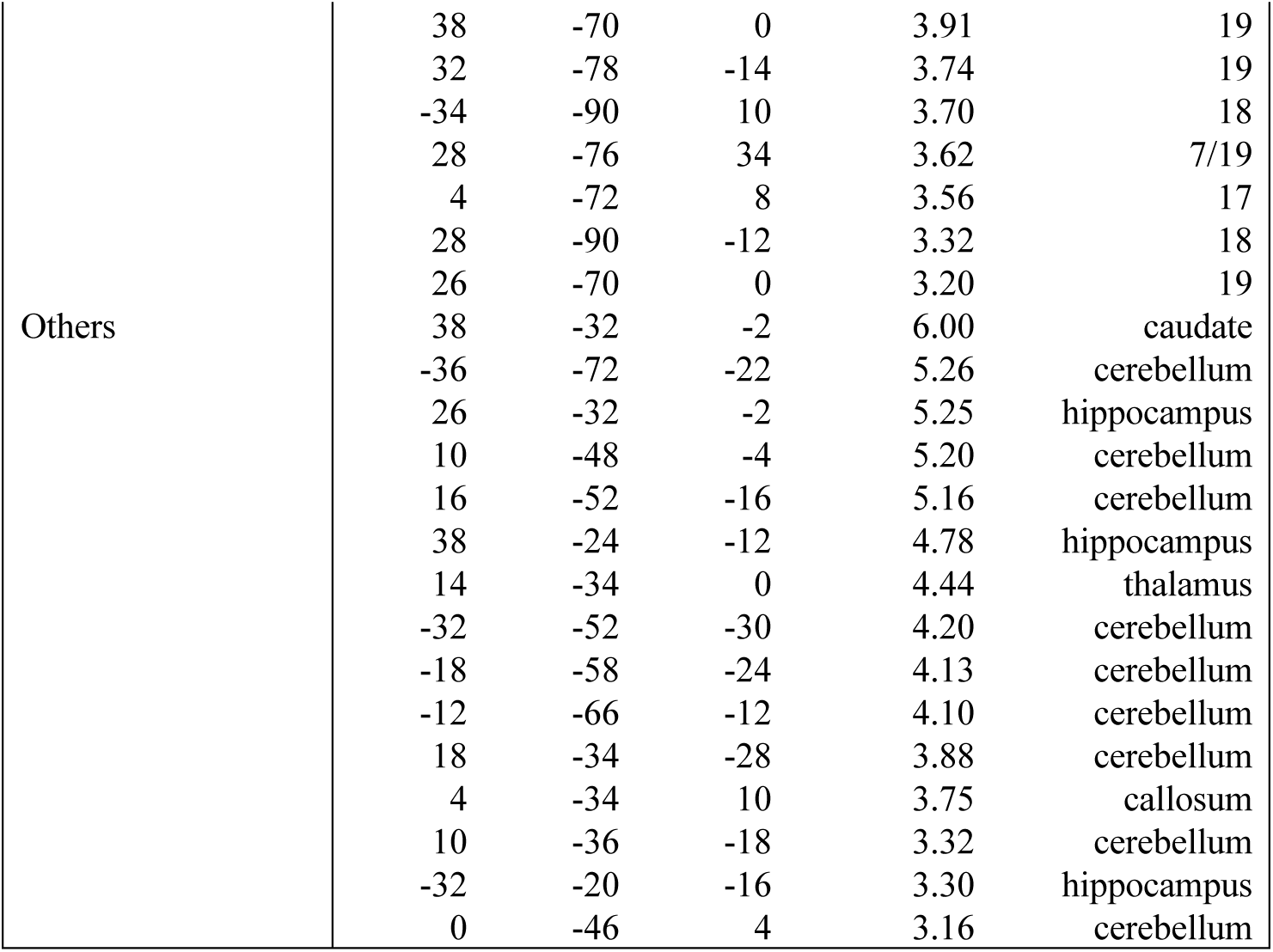
Brain regions showing significant accuracy to classify task dimension (face/place) in 40% coherence trials. Coordinates are listed in MNI space. Positive z-values indicate higher accuracy than chance level.

**Table S11.**
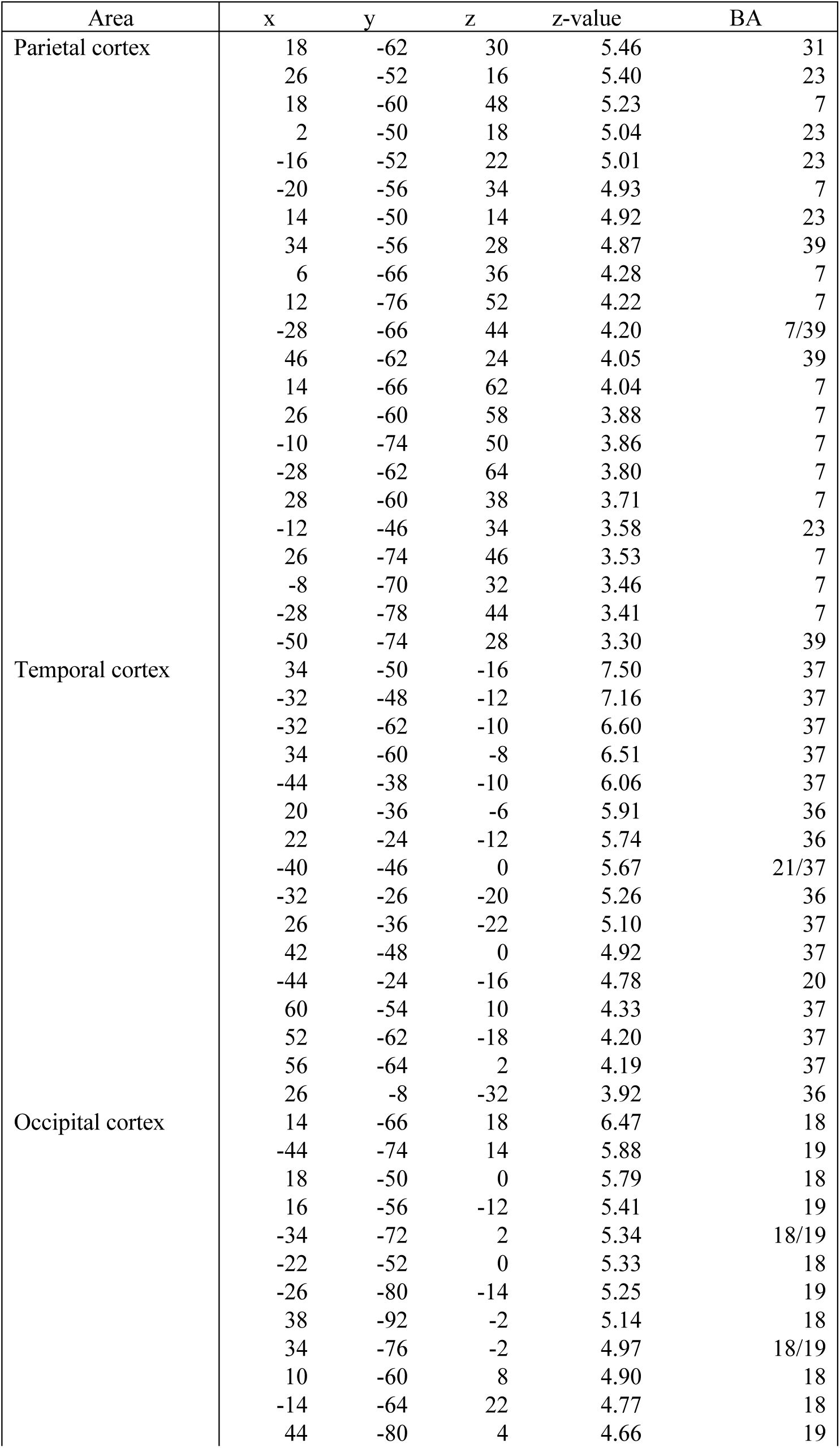
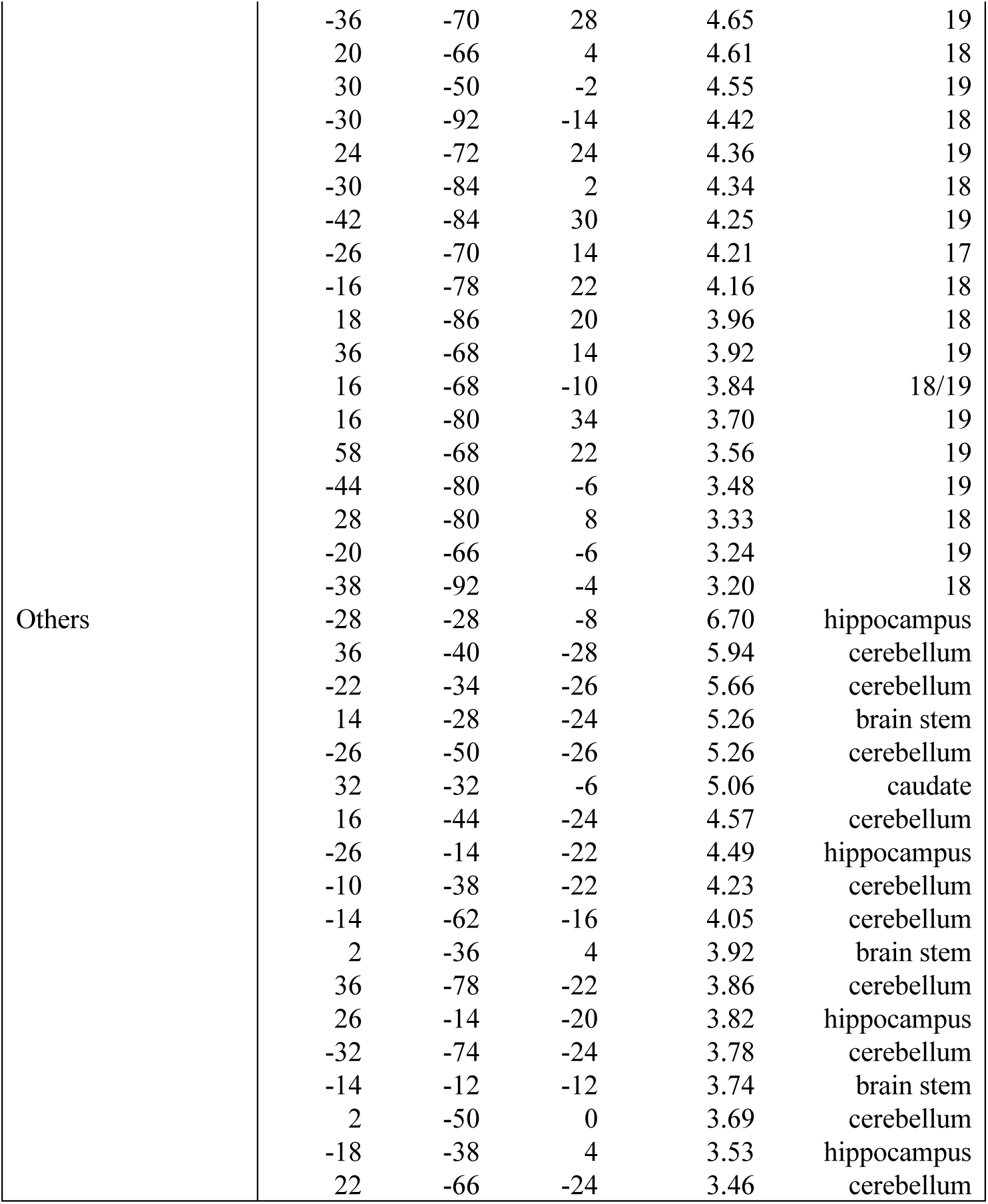
Brain regions showing significant accuracy to classify task dimension (face/place) in 80% coherence trials. Coordinates are listed in MNI space. Positive z-values indicate higher accuracy than chance level.

